# Both the low-density lipoprotein receptor and apolipoprotein E define blood-borne high-density lipoprotein entry into the brain

**DOI:** 10.1101/2025.05.23.655828

**Authors:** Sofia Kakava, Mary J. Gonzalez Melo, Wai Hang Cheng, Ilaria Del Gaudio, David Viertl, Stephanie Bernhard, Eveline Schlumpf, Mikaël Croyal, Evelina Voloviceva, Jianjia Fan, Carlos J. Barron, Antonis Katsoulas, Vasileia Kalaitzaki, Nora Jüdt, Thorsten Buch, Eric Camerer, Cheryl L. Wellington, Arnold von Eckardstein, Jerome Robert

## Abstract

Data from epidemiological and genetic studies as well as animal experiments indicate that high-density lipoproteins (HDL) play a role in the pathogenesis of central nervous system (CNS) diseases. Apolipoprotein A-I, the major protein of HDL, has been immunolocalized in the brain although it is produced exclusively by the liver and intestine. We therefore investigated how HDL cross the blood-brain barrier (BBB), using both *in vitro* and *in vivo* approaches. *In vitro*, we found that HDL bind to, are internalized by, and are transported through human brain endothelial cells via mechanisms involving the scavenger receptor BI (SR-BI) and the low-density lipoprotein receptor (LDLR). Notably, we discovered that LDLR facilitates only the transport of HDL particles containing apolipoprotein E (apoE). *In vivo,* HDL injected into the bloodstream enter into the brain through brain endothelial cells, and accumulated in medulla, cerebellum, olfactory bulb, hippocampus and cortex. Further investigation in *Ldlr^-/-^* mice revealed region-specific changes in HDL accumulation with reduced levels in the medulla and mixture of midbrain/cortex/hippocampus, no change in the olfactory bulb, and increased levels in the cerebellum. Together, these findings provide new insight on the interaction of the lipoprotein metabolism between the periphery and CNS. For the first time, we show that brain endothelial receptors and HDL composition jointly dictate HDL’s crossing through the BBB and their localization within the brain.

## Introduction

High-density lipoproteins (HDL) are lipid transporters within the bloodstream produced by the liver and intestines. HDL particles are a heterogeneous lipoprotein class that vary in size, shape, and composition with over 200 distinct proteins ^1^ and lipids ^2^. These do not appear uniformly across all HDL particles but instead form specialized subpopulations, each contributing to HDL’s vasoprotective, antithrombotic, antioxidant, anti-inflammatory, and cytoprotective functions ^3^. Initially investigated for their potential cardiovascular benefits ^4,5^, HDL have now been implicated in a wider range of conditions, including neurodegenerative diseases ^6^. Several studies indicate an inverse correlation between HDL-cholesterol (HDL-C) levels and dementia risk ^7–9^, although both low and very high HDL-C levels have shown associations with increased dementia risk ^10,11^. Recent large-scale genome-wide association studies (GWAS) on Alzheimer’s disease (AD) further underscore the link between HDL and brain health, identifying genes tied to HDL metabolism—such as *APOA1, APOE, ABCA1, APOC1, APOM, APOA2, PON1, CLU, LCAT*, and *CETP*—as significant contributors to AD risk ^12,13^. Notably, while certain HDL components, including apolipoprotein A-I (apoA-I), are present in the brain despite not being synthesized in the central nervous system (CNS) ^14^, HDL’s neuroprotective roles may depend on its ability to interact with and cross the blood-brain barrier (BBB). Current knowledge of HDL transport across endothelial cells is primarily derived from *in vitro* studies in aortic endothelium, where HDL trafficking is regulated by scavenger receptor BI (SR-BI) and ATP-binding cassette transporter G1 (ABCG1) ^15^. However, it remains unclear if these pathways are conserved in brain endothelial cells, which have limited trafficking capacity compared to aortic endothelial cells ^16–18^. Previously, we demonstrated that brain endothelial cells, unlike aortic endothelial cells, degrade rather than transport low-density lipoproteins (LDL), another lipid transporter in the bloodstream ^19^. Here, we investigate HDL transport across the BBB using both *in vitro* and *in vivo* approaches.

## Methods

### Animals

Animal experiments were conducted in strict accordance with ethical principles and guidelines for scientific experiments on animals set forth by the Swiss Academy of Medical Sciences. Protocols were approved by the ethics committee for animal experimentation of the canton of Vaud (VD-3781). Both male and female wild-type C57Bl/6NCrl mice (strain code: 632) were obtained from Charles River at the age of 7 weeks. The animals were housed in individually ventilated cages under controlled humidity and temperature conditions (21–23°C) on a 12-hour light/dark cycle and had access to food and water *ad libitum*.

To investigate the role of LDLR in the brain, animal procedures were conducted with approval from the Canadian Council on Animal Care and University of British Columbia (UBC, Vancouver, Canada) (A22-0181-A001). Male C57Bl/6J wild-type (IMSR_JAX:000664) and *Ldlr* knockout (B6.129S7-Ldlr ^tm1Her^ /J, IMSR_JAX:002207, refered as *Ldlr*^-/-^) mice were obtained from Jackson Laboratory at the age of 7 weeks. Animals were maintained on a chow diet (PMI LabDiet 5010; LabDiet) for 5 to 7 weeks and housed under controlled conditions with a 12-hour light/dark cycle, with *ad libitum* access to food and water.

For the vascular distribution, the study performed in Paris was conformed to the guidelines from Directive 2010/63/EU of the European Parliament on the protection of animals used for scientific purposes. Wild type C57BL/6J mice were purchased from Janviers lab. Mice were housed in a specific-pathogen free facility in a temperature-controlled environment, using a 12-hours light/dark cycle. Mice were fed a regular chow diet (A03-10, Scientific Animal Food and Engineering, Augy, France). Water and food were provided *ad libitum*.

### Cell culture

The human cortical microvascular cell line hCMEC/D3 (Sigma Aldrich, Switzerland; passage 34 to 38) and the primary human brain microvascular endothelial cells (hBMEC, Sciencell, USA; passage 4 to 7) were cultured in EBM-2 endothelial growth medium with BulletKit™ according to the manufacturer’s instructions (Cat. No. CC- 3156, Lonza, Switzerland). The medium was supplemented with 5% fetal bovine serum (FBS, Cat. No. 10500056, ThermoFisher Scientific, USA) and 5% heat-inactivated FBS (Cat. No. A4766801, Gibco, USA). The mouse brain endothelial cell line bEnd.3 (ATCC, USA; passages 8–10 after purchase) was cultured in Dulbecco’s Modified Eagle’s Medium (DMEM) (Cat. No. 12800-017, Sigma Aldrich, USA) containing high glucose, 1500 mg/L sodium bicarbonate, and 10% FBS. All cells were maintained in a humidified incubator at 37°C with 5% carbon dioxide.

### Lipoprotein isolation

LDL (1.019 < d < 1.063 kg/L) and HDL (1.063 < d < 1.21 kg/L) were isolated from the plasma of normolipidemic donors obtained from the Blutspende Zurich. Lipoproteins were isolated from apoE3/E3 individuals by density gradient ultracentrifugation as previously described ^20^. Briefly, plasma was thawed, and EDTA was added to a final concentration of 30 mM. The density was first adjusted to 1.069 kg/L using potassium bromide (Cat. No. 793604, Sigma-Aldrich, USA) to separate the HDL and LDL fractions. After centrifugation, the upper fraction containing LDL was collected, and the density of the lower fraction was adjusted to 1.21 kg/L for the isolation of HDL. Following a subsequent centrifugation, the upper fraction containing HDL was collected. Both HDL and LDL fractions were extensively dialyzed against 150 mM NaCl and 0.4 mM EDTA (pH 7.4) at 4°C. All centrifugations were performed at 59,000 rpm at 15°C for at least 16 hours using an Optima™ L-90K Ultracentrifuge with Type 70 Ti or Type 90 Ti fixed-angle rotors (Beckman Coulter, USA). Density was confirmed using a DS7000 densitometer (Kruess, Germany). HDL purity was assessed by sodium dodecyl sulfate-polyacrylamide gel electrophoresis (SDS-PAGE) followed by Coomassie blue staining.

HDL was further fractionated into particles containing apoE (HDLE+) and lacking apoE (HDLE-) using immunoaffinity chromatography. Briefly, HDL was loaded onto a 1 mL human anti-apoE resin column (Goat anti-Human Apo E [G05] Agarose immobilized, Cat. No. S81-118, Fortis Life Sciences, USA) at a concentration of 23.5 mg total protein/mL of PBS. After overnight incubation at 4°C with constant rotation (13 rpm), the flow-through (HDLE-) was collected with two 1 mL PBS washes. The bound HDLE+ particles were eluted after three incubations with 1 mL of 3 M sodium thiocyanate (NaSCN, Cat. No. 251410, Sigma-Aldrich, USA) followed by two 1 mL PBS washes. HDLE- particles were also incubated with NaSCN at the same protein-to-NaSCN ratio. Both fractions were extensively dialyzed against 150 mM NaCl and 0.4 mM EDTA (pH 7.4) at 4°C and concentrated using Amicon Ultra 30K centrifugal filters (Cat. No. UFC503096, Merk Millipore, USA) following the manufacturer’s instructions. The presence or absence of apoE was confirmed by western blotting.

### ApoE isoform phenotyping

ApoE isoforms were identified from 50 µL of plasma using a validated multiplexed assay involving trypsin proteolysis followed by the analysis of proteotypic peptides via liquid chromatography-tandem mass spectrometry, as previously described ^21–24^.

### Lipoprotein labelling

HDL or LDL were radiolabeled with Na^125^I (Hartmann Analytic, Germany) following the McFarlane monochloride procedure, modified for lipoproteins ^15,25^. Briefly, lipoproteins were diluted with physiological NaCl solution to a concentration of 37.5 mg total protein/mL. Immediately after adding glycine-NaOH at a concentration of 0.4 M, 37 MBq of ^125^I in 40 mM NaOH was added to the mixture. Labeling was initiated by adding HCl (0.17 M), ICl (8 mM), and NaCl (1.6 M) at a ratio of 1:4. After 5 minutes of incubation at room temperature, unbound ^125^I was removed using a PD10 desalting column (Cat. No. 17085101, Cytiva, USA). Labeled lipoproteins were then extensively dialyzed against 150 mM NaCl and 0.4 mM EDTA (pH 7.4) at 4°C. Protein concentration was measured using a BCA Protein Assay Kit (Cat. No. UP95424A, UP95425A, Interchim, France), and the activity of radiolabeled material was measured using a Wizard2 γ-counter (PerkinElmer, USA).

HDL and bovine serum albumin (BSA) were fluorescently labeled on the protein moiety using Atto 655 NHS (Cat. No. AD655-35, ATTO-TEC GmbH, Germany) following the manufacturer’s instructions. For *in vivo* experiments, Atto655-HDL and Atto655-BSA were concentrated with Amicon Ultra 30k centrifuge filters to 20.4 mg/ml and 25 mg/ml respectively. For lipid moiety labeling, 1,1’-dioctadecyl-3,3,3’,3’- tetramethylindocarbocyanine perchlorate (DiI, Cat. No. D282, Invitrogen, USA) was used according to St. Clair’s protocol ^26^. Briefly, DiI was dissolved in DMSO to a final concentration of 3 mg/mL. Then, 5 mg of HDL was mixed with 10.5 mL of human lipoprotein-deficient serum (LPDS) and 250 μL of Dil in DMSO. After overnight incubation at 37°C in the dark, DiI-HDL was isolated by KBr gradient ultracentrifugation after adjusting the density to 1.21 g/L. The isolated DiI-HDL was extensively dialyzed against 0.4 mM EDTA and 150 mM NaCl (pH 7.4) at 4°C, protected from light. For particles labeled on both lipid and protein moieties, 1 mg of DiI-HDL was further labeled with Atto 655 NHS following the manufacturer’s instructions.

### HDL biodistribution in the mouse brain

Four mice (2 females and 2 males) were injected with 100 mg/kg of ^125^I-HDL into the tail vein. Four hours after the injection, the animals were perfused, and the brains were collected for further analysis. The olfactory bulb, medulla, cerebellum, cortex, hippocampus, and the remaining brain tissue (referred to as rest of the brain, RB) were dissected, and the radioactivity was measured using a Wizard3” γ-counter (PerkinElmer, USA), Counts per minute (cpm) were normalized to the weight of each piece.

### Tail vein injection with fluorescently labelled HDL

Atto655-HDL, Atto655-BSA, or PBS (vehicle control) were injected into the tail vein of wild-type or *Ldlr*^-/-^ mice at a concentration of 100 mg/kg. All injections were performed between 8:00 and 11:00 AM to account for the circadian rhythm of lipoprotein metabolism ^27,28^. After injection, mice were returned to their cages with food and hydration gels provided. Brains were collected 4 hours post-injection.

EDTA blood was collected via cardiac puncture, centrifuged at 2,000g for 10 minutes at room temperature, and plasma was stored at -80°C. Animals were perfused for 7 minutes at a flow rate of 8 mL/min with PBS containing 2,500 U/L heparin.

For the pilot experiment, brain tissues were removed, divided into two hemispheres, and processed accordingly. One half of the brain was snap-frozen at -80°C. Frozen tissue was dissected to isolate the olfactory bulb, cerebellum, medulla, and the remaining brain tissue (referred to as RB) for human apoA-I quantification using ELISA. The other half of the brain was fixed in 4% paraformaldehyde (PFA) in PBS for 16 hours, followed by cryoprotection in gradient sucrose solutions (12%, 18%, 30%, each for approximately 16 hours) prior to cryostat sectioning for histological analysis. Based on this pilot experiment, a power calculation was performed for a two-sided test with α = 0.05 and 80% power, using the means and standard deviations of the ELISA quantification for midbrain. This analysis determined that 15 animals were required for subsequent experiments, which were performed identically as the pilot experiment.

### Histology

Brains injected with Atto655-HDL and Atto655-BSA were embedded in optimal cutting temperature (OCT) media and sectioned in the sagittal plane at a thickness of 25 μm using a cryostat (Leica, Germany). Sections were rehydrated in PBS and stained with 1 μg/mL of DAPI for 30 minutes before mounting. Slides were imaged using a ZEISS Axio Scan.Z1 Slide Scanner (Germany) or a Leica SP8 Inverted Confocal Microscope (Germany).

### Thick brain section

Atto655-HDL were administered via retro-orbital injection in wild type mice. After 4 hours, fresh brain tissue from transcardially perfused mice ((DPBS (Gibco) containing 2 U heparin (Sanofi Aventis France), followed by perfusion with 1% PFA diluted in DPBS with 2 U/mL heparin)) was sectioned into 0.8 mm slices using a tissue chopper (The Mickle Laboratory Engineering Co. Ltd, UK) and post-fixed in 4% PFA for 1 hour at room temperature. The sections were then blocked overnight at 4°C in TNBT buffer (0.1 M Tris, pH 7.4; 150 mM NaCl; 0.5% blocking reagent from Perkin Elmer; 0.5% Triton X-100) and washed with TNT buffer (0.1 M Tris, pH 7.4; 150 mM NaCl; 0.5% Triton X-100). Primary (mouse anti-αSMA-Cy3 1:250, Cat. No. C6198, Sigma; goat anti-CD31 1:200, Cat. No. AF3628, R&D Systems) and secondary antibodies (donkey anti-primary antibody coupled with AlexaFluor488 1:400, Thermofisher) were incubated overnight at 4°C. Finally, the thick sections were mounted in Dako Fluorescence Mounting Medium (Cat No. S3023, Dako). Confocal images were acquired on an inverted Leica SP8 laser scanning microscope using LAS-X software. For image acquisition, 63x (NA 1.40) oil immersion objective was used

### Blood Vessel Isolation and Immunostaining

Mouse brain vessels were isolated as previously described ^29^ with the following modifications. Briefly, brains from three wild-type mice injected with Atto655-HDL were pooled and kept frozen until processing. Brains were finely minced and homogenized using a Dounce homogenizer in ice-cold PBS containing 1% BSA. The homogenate were centrifuged at 2,000 × g for 10 minutes at 4°C to pellet the vessels, followed by resuspension in a 17.5% dextran solution and centrifugation at 4,400 × g for 15 minutes at 4°C to separate vessels from myelin and debris. The vessel pellet was resuspended in PBS with 1% BSA and passed through a 100 µm cell strainer to obtain purified brain microvessels for immunostaining.

Isolated vessels were transferred to 0.2 mL tubes and allowed to settle for 1 hour at 25°C before fixation with 4% PFA for 5 minutes. Vessels were then blocked with 1% BSA and 5% normal donkey serum for 1 hour at room temperature (RT), followed by overnight incubation at 4°C with primary anti-CD31 antibody (Abcam, Cat. No. ab7388, 1:200 dilution). The next day, samples were incubated with Alexa Fluor 488-conjugated donkey anti-rat secondary antibody (Invitrogen, Cat. No. A48269, 1:500 dilution) for 1 hour at RT. After washing, nuclei were stained with DAPI (Sigma Aldrich, cat No. 62248, 1 μg/ml) for 5 minutes. Samples were mounted on glass slides using Fluorsave mounting medium (Merck Millipore, Germany), and images were acquired using a Leica SP8 inverted confocal microscope.

### Lipoprotein cell binding, association and transport

For binding and association assays, cells were seeded at a density of 1.2 × 10^5^ cells per well in a 24-well plate and cultured for 72 hours until confluent. For binding assays, cells were washed with ice-cold DMEM containing 25 mM HEPES (Cat. No. 12430- 054, Gibco, Switzerland) and 0.2% BSA. Cells were then placed on ice in the same medium. After 30 minutes, cells were incubated with ice-cold DMEM (25 mM HEPES and 0.2% BSA) containing 10–20 μg/mL of ^125^I-HDL, ^125^I-HDLE+, ^125^I-HDLE-, or ^125^I-LDL in the absence (total binding) or presence (non-specific binding) of a 40-fold excess of unlabeled HDL, LDL, or BSA. After 1 hour, cells were washed twice with 1 mL of ice-cold Tris-BSA buffer (50 mM Tris, 150 mM NaCl, 0.02% NaN3, and 0.2% BSA) and once with ice cold PBS containing 1 mM MgCl2 and 0.1 mM CaCl2, before being lysed with 0.1 M NaOH. Lysed cells were then counted using a Wizard2 γ-counter (PerkinElmer, USA). Cpm values were normalized to the activity of the respective iodination and to the total protein content of the cells, measured using the DC Protein Assay (Cat. No. 5000112, Bio-Rad Laboratories, USA) following the manufacturer’s instructions.

For association assays, cells were incubated at 37°C for 1 hour with DMEM (25 mM HEPES and 0.2% BSA) containing 10–20 μg/mL of ^125^I-HDL, ^125^I-HDLE+, ^125^I-HDLE-, or ^125^I-LDL in the absence (total association) or presence (non-specific association) of a 40× excess of unlabeled HDL, LDL, or BSA. For time-course experiments, cells were incubated under the same conditions for the indicated durations. After incubation, cells were processed as described for the binding assay.

For transport assays, endothelial cells were seeded at a density of 1 × 10^5^ cells per well in a 24-well plate on 0.4 μm pore-size PET membrane inserts (Cat. No. 353095, Corning, USA) that had been pre-coated with 50 μg/mL of Collagen I (Cat. No. A10483- 01, Gibco, Thermo Fisher Scientific, USA for 1 hour at room temperature. After 3 days, once the cells had grown to confluence, they were incubated with 20 μg/mL of ^125^I- HDL, ^125^I-HDLE+, or ^125^I-HDLE-in the absence (total transport) or presence (non-specific transport) of a 60× excess of unlabeled HDL or LDL at 37°C. After 1 hour, media in the bottom chamber was collected and counted as for the binding assay to determine the amount of transported lipoproteins.

Specific binding, association, and transport were calculated by subtracting the normalized cpm values of the non-specific condition from the normalized values of the total condition.

### HDL degradation

To measure HDL degradation, endothelial cells were seeded at a density of 1 × 10^5^ cells per well in 24-well plates and cultured to confluence over 3 days. On the day of the assay, cells were incubated as described above for HDL association, with 10 μg/mL of ^125^I-HDL in DMEM (25 mM HEPES and 0.2% BSA) in the absence (total degradation) or presence (non-specific degradation) of a 40× excess of unlabeled HDL. After 4 hours of incubation, the assay media was collected and mixed with ice-cold trichloroacetic acid (TCA, Sigma Aldrich, Switzerland) to a final concentration of 12%. Samples were incubated on ice for 30 minutes at 4°C and then centrifuged at 2000× g for 10 minutes at 4°C. Supernatants were transferred to new tubes containing NaI (final concentration 0.4%), vortexed, and incubated for 5 minutes at room temperature. H2O2 was then added to a final concentration of 1.1%. After mixing and 5 minutes of incubation, degradation products were isolated using chloroform extraction. The upper aqueous phase was collected and counted using a Wizard2 γ- counter. In parallel, cells were processed as described for the HDL association assay. The percentage of degradation relative to association was calculated by dividing the cpm of degradation products by the sum of the cpm of association and degradation products, multiplied by 100.

### HDL internalization

Endothelial cells were seeded at a density of 3 × 10^5^ cells per well in a 12-well plate and grown to confluence over 3 days. Cells were then incubated with 50 μg/mL of Atto655-HDLE+ or Atto655-HDLE-, with or without a 100× excess of unlabeled HDL (non-specific competitor), in DMEM containing 25 mM HEPES and 0.2% BSA. After 3 hours of incubation, cells were detached by incubating with Accutase (Cat. No. A6964, Sigma-Aldrich, USA) for 5 minutes at 37°C. Detached cells were collected and washed twice with ice-cold PBS. To exclude signals from dead cells, samples were incubated with Propidium Iodide (PI) (Cat. No. 81845, Fluka, Switzerland) at a final concentration of 1 μg/mL and FITC-Annexin V (Cat. No. 640945, BioLegend, USA) at a final concentration of 2 μg/mL. The uptake of fluorescent material by live cells was measured using a BD LSR II Fortessa flow cytometer (BD Biosciences, USA) and analyzed with BD FACSDIVA™ software. During acquisition, forward scatter versus side scatter gating (FSC-A/SSC-A) was applied to exclude cell debris, followed by gating to exclude doublets (FSC-A/FSC-H). Double-negative cells for both FITC-Annexin V and PI were selected by gating (FITC-A/PI-A). To account for spectral overlap between fluorophores, unstained and single-stained controls were acquired and analyzed using the compensation function in the FACSDIVA™ software. For all experiments, approximately 10^4^ events per condition were recorded within the final gate containing the live cell population and used for analysis. Data were analyzed using FlowJo Software (v10.10) (FlowJo LLC, USA). The median fluorescence intensity (MFI) of each population was used to compare the different conditions, and specific uptake was calculated as the difference between total uptake and non-specific uptake for each condition.

### Size-exclusion chromatography

The particle size of HDL, before or after transport, was determined using Fast Protein Liquid Chromatography (FPLC) (ÄKTA Go, Cytiva, USA). Briefly, 72 hours after seeding endothelial cells on transwells as described above for transport assays, 30 μg/mL of ^125^I-HDL, ^125^I-HDLE+ or ^125^I-HDLE- were added to the top chamber and incubated for 1 hour at 37°C. Media from the bottom chamber was collected and mixed with saturated sucrose (ratio 2:1) and 1 mg of non-labeled HDL to stabilize the particles. For size separation, 500 μL of the collected material was applied to a HiLoad 16/600 Superdex 200pg size exclusion column (Cat. No. 28989335, Cytiva, USA). Isocratic elution was performed at 4°C using a buffer containing 150 mM NaCl and 10 mM Tris, pH 7.5, at a constant flow rate of 1 mL/min. Fractionation began immediately after injection, with fractions of 1 mL each collected. Each fraction was measured using a gamma counter to determine the distribution of radioiodinated particles transported across the endothelium. For all experiments, transwells without cells served as a negative control to assess spontaneous modifications of particle size.

### RNA interference

Cells were seeded and reverse transfected using specific siRNA targeting selected transcripts. Specifically, siRNA against *LDLR* (SMARTPool, Dharmacon, USA, L- 011073-00-0010), *SCARB1* (Ambion, USA, numbers s2648 #4390824 and s2649 #4390825), *ACVRL1* (SMARTPool, Dharmacon, D-005302-06-0010 or Ambion #4392420), *CAV1* (SMARTPool, Dharmacon, #M-003467-01-0005), *AP2M1* (SMARTPool, Dharmacon, #L-008170-00-0005), *LRP1* (SMARTPool, Dharmacon, #M-004721-00-05), *LRP8* (SMARTPool, Dharmacon, #M-011902-01-0005), *ABCG1* (Dharmacon, #L-008615-00), and respective non-targeting scramble siRNA (Dharmacon SMARTPool, #D-001810-10-50 or Ambion #4390843 or #4390846) were diluted in Opti-MEM (Cat. No. 31985-070, Gibco, USA).

Lipofectamine RNAiMAX transfection reagent (Cat. No. 13778150, ThermoFisher Scientific, USA) was diluted in Opti-MEM, and the two solutions were mixed at a 1:1 ratio. After 30 minutes of incubation at room temperature, 100 μL of the siRNA/lipofectamine working solution was distributed per well of a 24-well plate. hCMEC/D3 or hBMEC cells were then seeded in 400 μL of complete EGM-2 medium containing 10% FBS at a density of 1 × 10⁵ cells/well and cultured for 3 days until confluence. The final concentration of each siRNA and lipofectamine after the addition of the cells was 20 nM and 1.3 μl/mL respectively. For transfected cells in the transwell system, the media was changed 8–16 hours after transfection.

### LDLR and SR-BI pharmacological inhibition

Protein interference against SR-BI and LDLR was achieved using a 30-minute pre-treatment with a blocking antibody against SR-BI (Cat. No. NB400-131, Novus Biological, RRID: AB_10002812, 1:200) or LDLR (Cat. No. sc18823 [C7], Santa Cruz Biotechnology, USA, RRID: AB_627881, 1:50), goat IgG control (Cat. No. ab37373, Abcam, USA, RRID: AB_3665323, 1:200), or mouse IgG2b control (Cat. No. sc-3879, Santa Cruz Biotechnology, USA, RRID: AB_737262, 1:200) in DMEM containing 25 mM HEPES and 0.2% BSA. The association assay with ^125^I-HDL was then performed as described above. Antibodies were maintained in the media throughout the assay. To block heparin sulfate proteoglycans (HSPG), cells were pre-incubated for 1 hour with 0.2 mg/mL of heparin (Cat. No. H2149, Sigma-Aldrich, USA followed by association assay with ^125^I-HDL as described above. Heparin was maintained in the media throughout the assay.

### Quantitative Reverse transcriptase PCR

Total RNA was isolated using the RNA isolation kit (Cat. No. 740955, Macherey Nagel, Germany, NucleoSpin™ Mini Kit for RNA Purification) and cDNAs were generated using the RevertAid First Strand Synthesis kit (Cat. No. K1621, Thermo Fisher Scientific, USA) according to manufacturer’s instructions. Real-time quantitative PCR reactions were carried out on a Roche Light Cycler 480-II (Roche, Switzerland) using the LightCycler® 480 SYBR Green I Master Mix (Cat. No. 04887352001, Roche, Switzerland) and specific primers as shown in the table below

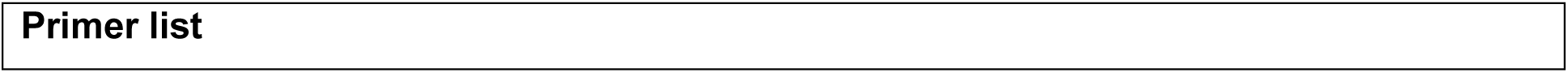
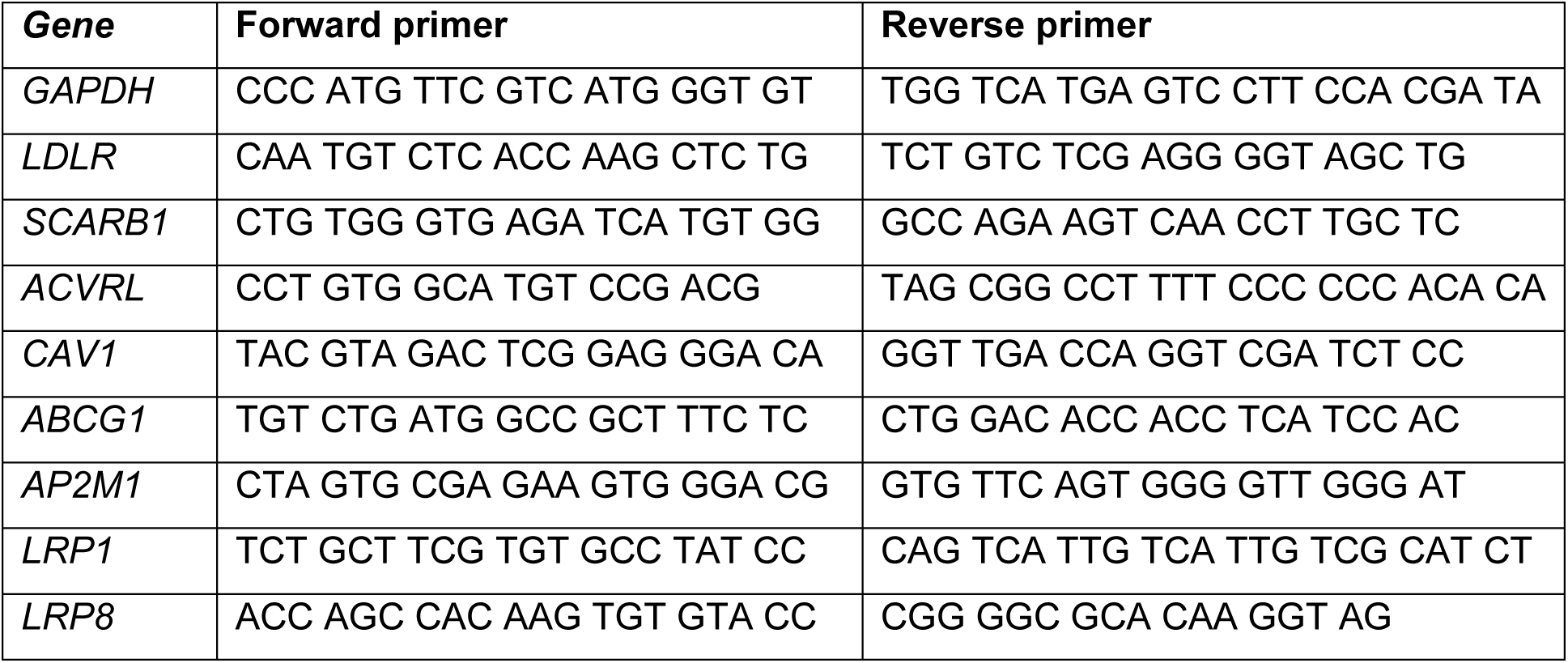

### SDS PAGE and Western Blot

All HDL fractions were quantified using the DC assay. Twenty-five μg of the original HDL pool and each isolated subpopulation, HDLE+ and HDLE-, were separated by SDS-PAGE followed by electrophoretic transfer to polyvinylidene fluoride (PVDF) membranes (GE Healthcare, USA). Membranes were first blocked for 1 hour with 5% skim milk powder solubilized in PBS containing 0.1% Tween (PBST), then incubated with antibodies against apoE (Cat. No. ab1906, Abcam, USA, RRID: AB_302668, 1:1000) or apoAI (Cat. No. 600-101-109, Rockland, USA, RRID: AB_2056545, 1:1000). After 1 hour at room temperature or overnight at 4°C, membranes were extensively washed with PBST and incubated with secondary anti-goat or anti-mouse antibody (Agilent Dako, USA, 1:1000) in blocking buffer. After 1 h, membranes were washed extensively in PBST and developed using SuperSignal^TM^ West Pico or Femto PLUS chemiluminescence substrate (ThermoFisher Scientific, USA) with a Fusion FX imager (Agilent, Switzerland).

Brain tissues were lysed in RIPA buffer and centrifuged for 15 minutes at 20,000 g to remove any cell debris. Supernatant was collected and proteins were quantified using the DC assay. Twenty-five μg of total protein were separated by SDS-PAGE followed by electrophoretic transfer to PVDF membranes. Atto655-HDL or Atto655-BSA were detected by measuring the fluorescence at the 655 nm channel using the ChemiDoc Imaging System (Bio-Rad Laboratories, USA). Densitometry of the images was captured with ImageJ (NIH, USA) and band intensity normalized to beta-actin (Cat. No. A5441, clone AC-15, Sigma-Aldrich, USA, RRID:AB_476744, 1:2500) as a loading control.

### ELISA quantification of human apoA-I

HDL levels in the mouse brain were quantified using human apoA-I ELISA (ab108803, Abcam, USA) following manufacturer’s instructions. Briefly, brain regions were weighted and lysed using 1 ml/mg of lysis buffer containing 10x PBS,1% Triton-X-100 and protease inhibitor cocktail (1 tablet per 10 ml, cOmplete Mini, Cat. No. 11836153001, Roche, Switzerland). Lysates were run through a 21G syringe to disrupt the tissue and further homogenized for 2x 3 min at 4°C and 20 Hz using a Tissue Lyser II system (QIAGEN, Germany). Sample were then sonicated for 30 seconds at 4°C with 10% magnitude using Digital Sonifier 450 tip sonicator (Branson Ultrasonics, Germany) and centrifuged for 15 minutes at 10°C and 14.000g. The supernatant was transferred and aliquoted to new tubes and stored at 80°C until quantification. The mean absorbance value of BSA injected samples were subtracted to the values of HDL injected mice and apoA-I concentration was normalized to total protein concentration.

### Microscopy

Cells were seeded at a density of 1 × 10^5^ cells per glass coverslip (13 mm round, Cat. No. SUP0117530, Southern Cross Science, Australia) in a 24-well plate and grown for 3 days until confluence. On the day of the assay, cells were incubated with 50 μg/ml of fluorescently labeled HDL, HDLE+, HDLE-, or double-labeled HDL for 1, 4, or 24 hours in DMEM containing 25 mM HEPES and 0.2% BSA without FBS. Cells were then washed twice with ice-cold PBS and fixed with 4% PFA for 15-20 minutes. Nuclei were counterstained with DAPI in PBS (1 μg/ml) for 30 minutes before mounting coverslips on positive-charge slides using ProLong Gold Antifade Reagent (Cat. No. P36934, Invitrogen, USA). Lysosomes were stained after incubation with HDL by adding 50 nM Lysotracker (LysoTracker Red DND99, Cat. No. L7528, Invitrogen ThermoFisher Scientific, USA) in DMEM without serum for 15 minutes at room temperature, followed by PBS washes and DAPI counterstaining. Cells were imaged using a Leica Inverted SP8 or ZEISS LSM 900 confocal microscope.

### Statistical analysis

For all in vitro experiments with the exception of RT-qPCR, linear raw data were first log transformed and then analyzed by a blocked (“experiment” and “treatment”) Student’s t-Test or one-way ANOVA with Dunnett’s post hoc test with “Experiment” being the blocking factor. For qRT-PCR analysis, 2^−ΔΔCt^ values were used in the same test. Data were obtained from at least three independent experiments and graphically represented as mean ± standard deviation (SD). For all *in vivo* experiments, raw data was first tested for normality using Shapiro-Wilk test followed by a parametric t-test. All statistical analyses were performed using SPSS statistic 25 (IBM, SPSS Inc., Chicago, IL, USA) and p-values < 0.05 were considered significant. Prism 8 (GraphPad Software, USA) was used to plot all data with the control condition normalized to 100%.

## Results

### 1. HDL enter the brain through endothelial cells and are found in different brain regions

To investigate the entry of blood-borne HDL into the CNS, we labelled HDL on the protein moiety with the far-red fluorescent dye Atto655 NSH ester. Four hours after intravenously injecting Atto655-HDL in the tail veins of wild-type mice, we extensively perfused them and isolated brains for further analyses (***Figure 1A***). Protein separation of brain lysates by SDS-PAGE revealed a fluorescent signal with an apparent molecular mass of around 29 kDa expected for apoA-I, the major constituent of HDL (***Figure 1B***) in the brain of HDL-injected mice but not in PBS injected control animals. Using microscopy, we found fluorescently-labeled plasma HDL in multiple brain regions, including the olfactory bulb (OB), cortex (CX), choroid plexus (CP), hippocampus (HP), thalamus (TH), hypothalamus (HT), midbrain (MB), medulla (MD), and cerebellum (CB) (***Figure 1C***). We then investigated the distribution of ^125^I-HDL injected into the tail veins of male and female wild type mice in dissected brain regions. Radioactivity was present in all regions analyzed, and was highest in OB, MD, and CB (***Figure 1D*).** We further investigated whether HDL enters the CNS through the cerebrovascular endothelium. In thick brain sections of wild-type mice injected with Atto655-HDL, immunostaining to detect endothelial cells (CD31+, green) showed that HDL (magenta) are found in vesicular structure within the cells. HDL signal was notably higher in veins and capillaries relative to α-smooth muscle actin (αSMA, blue) positive arteries (***Figure 1E***). Brain vasculature isolation followed by confocal imaging confirmed HDL entry through endothelial cells, with HDL (magenta) found in perinuclear vesicular structures within endothelial cells (CD31+). Interestingly HDL were also found in perivascular cells (CD31- cells) (***Figure 1F***).

**Figure 1:**
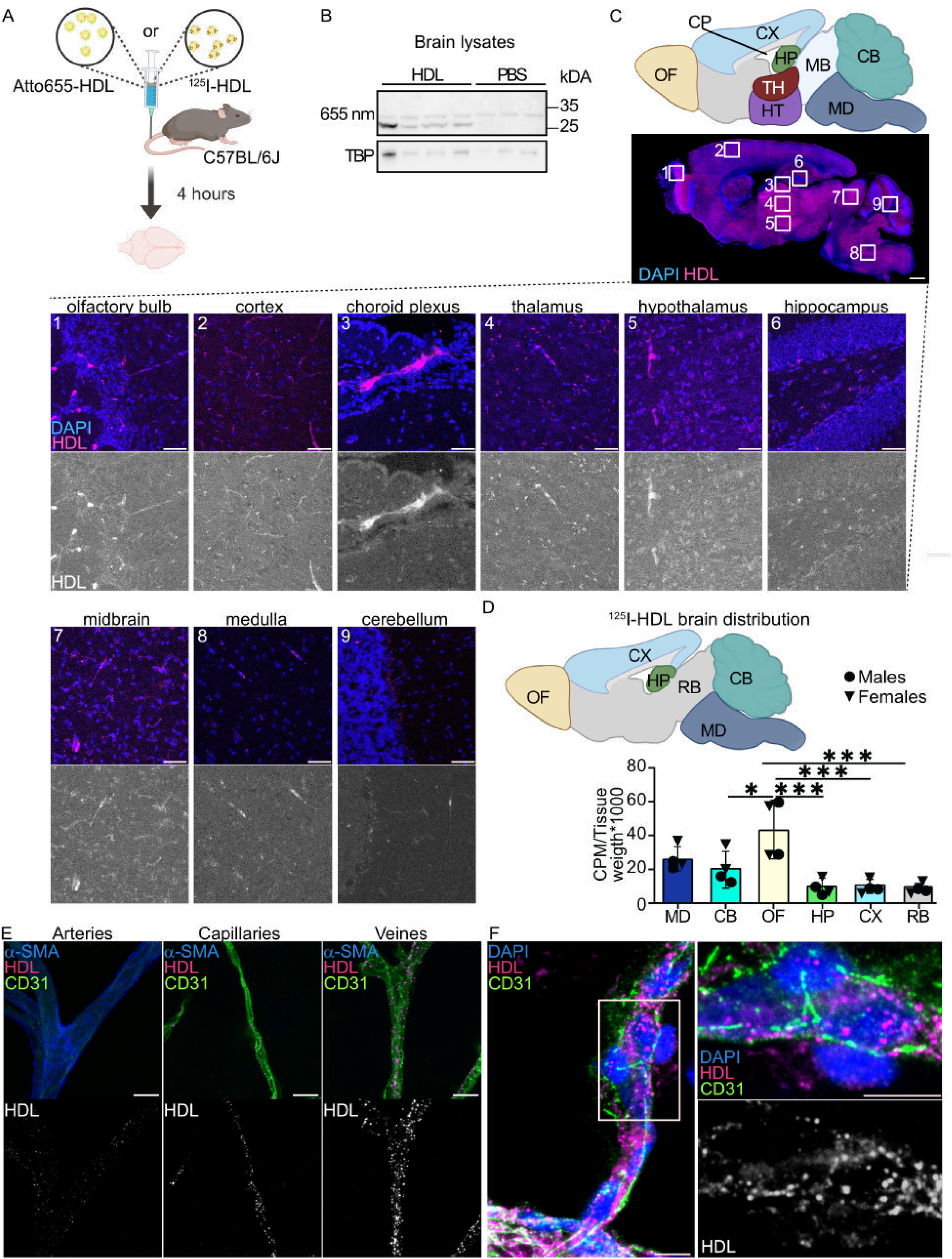
**Blood-borne HDL enters the brain through endothelial cells in a region- specific manner**. **A**) Schematic representation of the experiment, Atto655-HDL (100 mg/kg), or PBS vehicle control were injected in the tail veins of seven weeks old male wild-type mice. Four hours after injection, animals were transcardially perfused, and brains were collected, separated sagitally into two hemispheres for molecular assays and histology respectively. **B**) Brain lysates (25 μg) were separated by SDS-PAGE and atto655-HDL signal was recorded and compared to the loading control TATA-binding protein (TBP). **C**). Atto655-HDL (magenta) were visualized in 25 μm sagittal sections counterstained with DAPI and visualized using a slide scanner and confocal microscopes. Scale bar: 1000 μm for whole brain, 50 μm for zoomed-in images **D**) ^125^I-HDL (100 mg/kg) were injected into the tail veins of 7 weeks old male and female WT mice. After 4 hours and transcardiac perfusion, brains were dissected into medulla (MD), cerebellum (CB), olfactory bulb (OB), hippocampus (HP), cortex (CX) and remaining brain (RB) regions and ^125^I-HDL activity counted using a γ-counter. **F**) Fixed brains after Atto655-HDL (magenta) injection were sectioned in the coronal plane at 200 μm using vibratome. Vascular beds were visualized by immune staining against CD31 (green, endothelial cells) and αSMA (blue, arteries) and confocal microscopy. Scale bar: 10 μm. **G**) Brain vessels were isolated by gradient centrifugation, counterstained with DAPI and CD31 (green, endothelial cells) and imaged using confocal microscopy. Scale bar: 500 μm. Points in graphs represent individual animals, bars represent the mean and error bars ± SD: *p<0.05, **p < 0.01, ***p<0.001.

### 2. HDL uptake and transport through endothelial cells are regulated by SR-BI and LDLR

To gain deeper insights on HDL trafficking through brain endothelial cells, we incubated the brain endothelial cell line hCMEC/D3 with Atto655-HDL or HDL labeled on the lipid moiety (DiI-HDL). After one hour, both Atto655-HDL (***Figure 2A***) and DiI-HDL (***Figure 2B***) were found in perinuclear vesicles similar to the cerebrovascular pattern observed *in vivo* and that persisted up to 24 hours of incubation (***Supplemental Figure 1A-B***). Incubation of endothelial cells with HDL labeled on both the protein and lipid moiety showed that both Atto655 and DiI signals co-localized after one hour suggesting holoparticle uptake rather than as separate components (***Figure 2C***). HDL interaction with hCMEC/D3 was further characterized with respect to cellular association at 37°C, which corresponds to both binding and internalization, and transport using ^125^I-HDL. The association (***Figure 2D***) of ^125^I-HDL was significantly competed with a 40-fold excess of either non-labeled HDL or LDL but not albumin (BSA), suggesting a common pathway for both lipoproteins. By comparing the interactions of HDL with hCMEC/D3, primary human brain endothelial cells (hBMEC) (***Supplemental Figure 1C***), and the murine brain endothelial cells, Bend.3 (***Supplemental Figure 1D***), we found that HDL association is conserved in primary human cells and murine cells. We then measured the ability of hCMEC/D3 grown on transwell inserts to transport ^125^I-HDL from apical to basolateral chambers in the presence of excess non-labeled HDL and LDL. Albumin was not used as a competitor for these experiments as it did not compete for association. After one hour, only an excess of non-labeled HDL significantly reduced ^125^I-HDL (***Figure 2E***). Notably, gel filtration confirmed a similar Stokes’ diameter in the material recovered in the basolateral chamber and HDL not exposed to cells (***Figure 2F***). Together, these data suggest that HDL and LDL share common mechanisms or pathways for association but not transport and that HDL are transported as holoparticles by brain endothelial cells.

**Figure 2:**
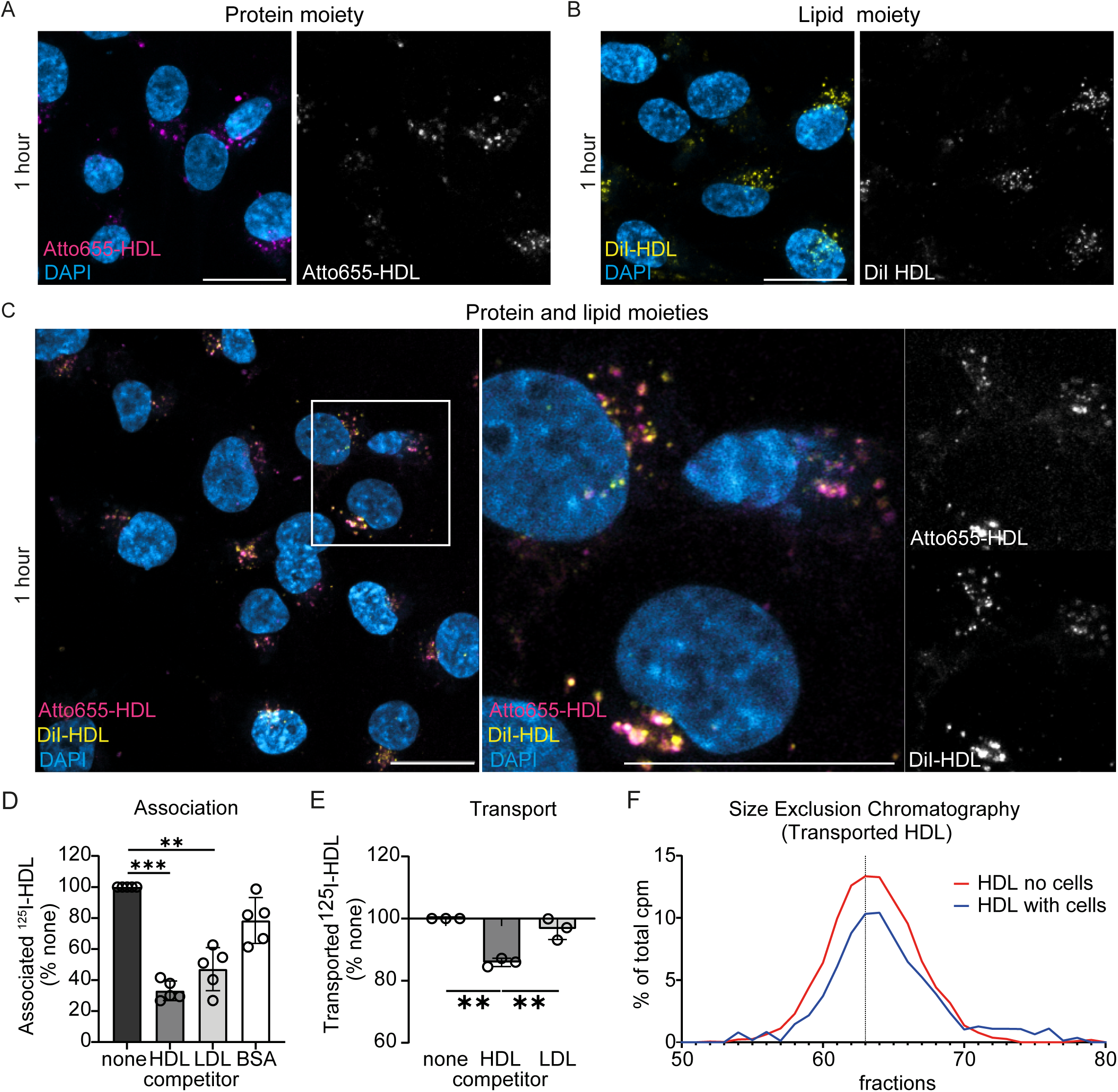
HDL are transported across human brain endothelial cells as holoparticles. HCMEC/D3 seeded on coverslips were incubated for 1 hour with 50 μg/ml of HDL labelled either on the protein moiety with atto655 NHS dye (**A,** magenta), on the lipid moiety using DiI (**B,** yellow) or on both moieties (**C**). **D**) Confluent hCMEC/D3 were incubated with 20 μg/ml of ^125^I-HDL without (none) or with 40× excess of non-labeled HDL, LDL, or BSA as competitors. After 1 hour, cells were washed, lysed and radioactivity was measured using a γ-counter **E**). Confluent hCMEC/D3 on transwells were incubated at 37°C with 20 μg/ml of ^125^I-HDL without (none) or with 40× excess of non-labeled HDL or LDL as competitor. After 1 hour, basolateral media was collected, and the radioactivity was measured using a γ-counter. **F**). Media from the basolateral chamber were loaded on a size exclusion column, 1 ml fractions were collected, and the radioactivity of each fraction was measured using a γ-counter. Red: ^125^I-HDL without cells contact, blue: ^125^I-HDL with cell contact. Representative image of 2 independent biological replicates. Points in graphs represent individual experiments (biological replicates, n = 2-4), bars represent the mean and error bars ± SD, *p<0.05, **p < 0.01, ***p<0.001.

To investigate transendothelial trafficking of HDL, we used specific siRNA’s to knock down expression of the known HDL receptors namely, scavenger receptor BI (SR-BI), and ATP binding cassette G1 (ABCG1) using specific siRNA. Because ^125^I-HDL association was also significantly reduced by an excess of non-labeled LDL and we previously showed that silencing SR-BI does not reduce LDL binding to brain endothelial cells ^19^, we also explored the role of the LDL receptor (LDLR) and activin A receptor like type 1 (ALK1), which regulate LDL uptake by endothelial cells ^19^. In publicly available databases of human and mouse single cell RNA sequencing (scRNASeq), we found that *SCARB1* (encoding SR-BI), *ABCG1*, *LDLR,* and *ACVRL1* (encoding ALK1) are expressed by brain endothelial cells ^30,31^. We also verified the expression of all receptors in our cultivated brain and aortic endothelial cells (***Supplemental Figure 2A***). We then measured knockdown efficiency by quantitative real-time PCR and found a significant down-regulation for each transcript (***Supplemental Figure 2B***). Finally, we investigated ^125^I-HDL association after silencing each receptor. In the absence of SR-BI, associated ^125^I-HDL was significantly reduced by 50% (***Figure 3A***) while the absence of ABCG1 (***Figure 3B***) and ALK1 (***Figure 3C***) did not significantly affect ^125^I-HDL association. Interestingly, we found that knocking down *LDLR* significantly reduced ^125^I-HDL association by 40% (***Figure 3D***). Knockdown of LDLR in primary human BMEC led to a similar significant reduction of both ^125^I-HDL and ^125^I-LDL association (***Supplemental Figure 2C***). Notably, knocking down *LDLR* did not alter the expression of *SCARB1* (***Supplemental Figure 2D***). To confirm the role of LDLR in HDL trafficking by brain endothelial cells, we overexpressed LDLR in hCMEC/D3 (***Supplemental Figure 2E***). In line with the silencing experiments, the association of ^125^I-HDL (***Figure 3E***) and ^125^I-LDL (***Supplemental Figure 2F***) were significantly increased in cells overexpressing LDLR while the expression of *SCARB1* remained unchanged (***Supplemental Figure 2G***). Because association integrates both binding and uptake of HDL, we used flow cytometry to measure specifically the uptake of HDL. Knockdown of LDLR led to a significant decrease in the internalization of Atto655-HDL (***Figure 3F***). Because we previously showed that LDL trafficking through LDLR in brain endothelial cells leads to particle degradation ^19^, we measured ^125^I-HDL degradation by hCMEC/D3 and primary hBMEC. After four hours, less than 5% of the associated HDL was degraded by either endothelial cell type (***Figure 3G***). Similarly, inhibition of clathrin-coated pit formation by silencing the AP-2 adaptor complex 1 (*AP2M1*) did not reduced ^125^I-HDL association (***Supplemental Figure 2H***). Inhibition of caveolin by silencing CAV1 also did not reduce ^125^I-HDL association (***Supplemental Figure 2I***). Finally, knocking down LDLR significantly reduced ^125^I-HDL transport through brain endothelial cells by 70% (***Figure 3H***). Together, these results suggest a novel role of LDLR in brain endothelial cells leading to HDL transport through endothelial cells but not to its degradation while internalized in endothelial cells.

**Figure 3:**
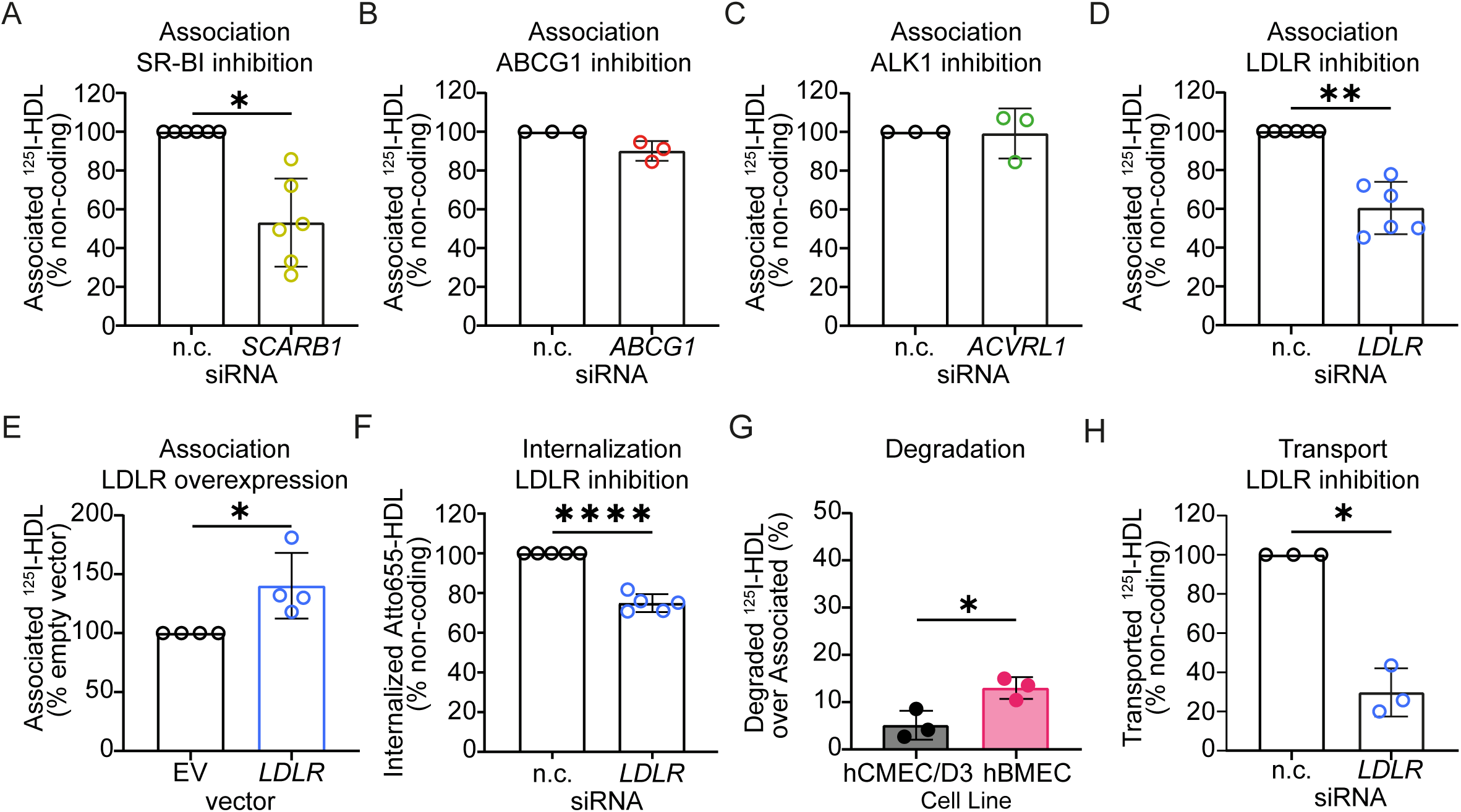
HDL transport through brain endothelial cells is reduced in the absence of LDLR. ^125^I-HDL association to hCMEC/D3 were measured as described in figure 2 72 hours after RNA interference against SCARB1 (**A**), ABCG1 (**B**), LDLR (**C**), and ACVRL1 (**D**) or transient overexpression of LDLR (**E**). **F**) Confluent hCMEC/D3 were incubated with 50 μg/ml Atto655-HDL. After 3 hours, cells were washed, detached and the median intensity fluorescence was quantified by flow cytometry. **G**) Confluent hCMEC/D3 and hBMEC were incubated with 10 μg/mL of ^125^I-HDL. After 4 hours, media was collected and degradation products were isolated after acid precipitation and chloroform extraction and compared to associated ^125^I-HDL. The percentage of degradation per association was calculated by dividing the CPM of degradation by the sum of association + degradation ×100. **H**) ^125^I-HDL association to hCMEC/D3 in the absence of clathrin complex (siRNA agains AP2M1) were measured as above **I**) The expression of LDLR was silenced in confluent hCMEC/D3 grown on transwell. ^125^I-HDL transport was measured as described in figure 2. Points in graphs represent individual experiments (biological replicates, n = 3–5), bars represent the mean and error bars ± SD, * p = 0.05 and ** p = 0.01.

### 3. LDLR preferentially regulates the trafficking of HDL containing apoE

One of the ligands of LDLR is apoE, which is present on 5-10% of blood-borne HDL particles ^32^. To investigate a possible interaction of HDL containing apoE (HDLE+) with LDLR in brain endothelial cells, we separated HDLE+ and HDL lacking apoE (HDLE-) using immunoaffinity chromatography. ApoE in both populations was assessed by SDS-PAGE and Western blotting. For the same apoA-I content, apoE was enhanced in HDLE+ compared to total HDL while apoE was not detectable in HDLE- (***Supplemental Figure 3A***). Because the release of HDLE+ from the immunoaffinity column required high-concentrated salts, we used gel filtration chromatography to confirm that HDL treated with NaSCN have a similar size distribution than native HDL (***Supplemental Figure 3B***). To compare HDLE+ and HDLE- trafficking in endothelial cells, both particles were radio- or fluorescently labeled on the protein moiety and association, internalization, and transport were measured. After 1 hour incubation, ^125^I- HDLE+ association was significantly higher than that of ^125^I-HDLE- (***Figure 4A***). Time course experiment showed enhanced association of HDLE+ over time compared to HDLE- with a higher saturation threshold (***Figure 4B***). Using Atto655-HDL and flow cytometry, we found that HDLE- uptake was also significantly lower compared to HDLE+ (***Figure 4C***). Although we showed that HDL are transported through and not degraded by brain endothelial cells, we also measured HDLE+ and HDLE- degradation, as in hepatocytes, the presence of apoE on HDL leads to particle degradation ^33^. After 4 hours, degradation of both HDLE+ and HDLE- was less than 5% of the respective associated HDL-subpopulation (***Supplemental Figure 3C***). Moreover, after 1 hour or 4 hours incubation neither of the two particles co-localized with the acidic marker lysotracker that labels lysosomes (***Supplemental Figure 3D***). Finally, the transport of HDLE- through brain endothelial cells cultured in transwells, was significantly lower compared to HDLE+ (***Figure 4D***). Notably, gel filtration chromatography revealed that HDLE+ particles have a larger Stokes’ diameter than HDLE- and both particle sizes remained similar after transport through brain endothelial cells (***Supplemental Figure 3E***).

**Figure 4:**
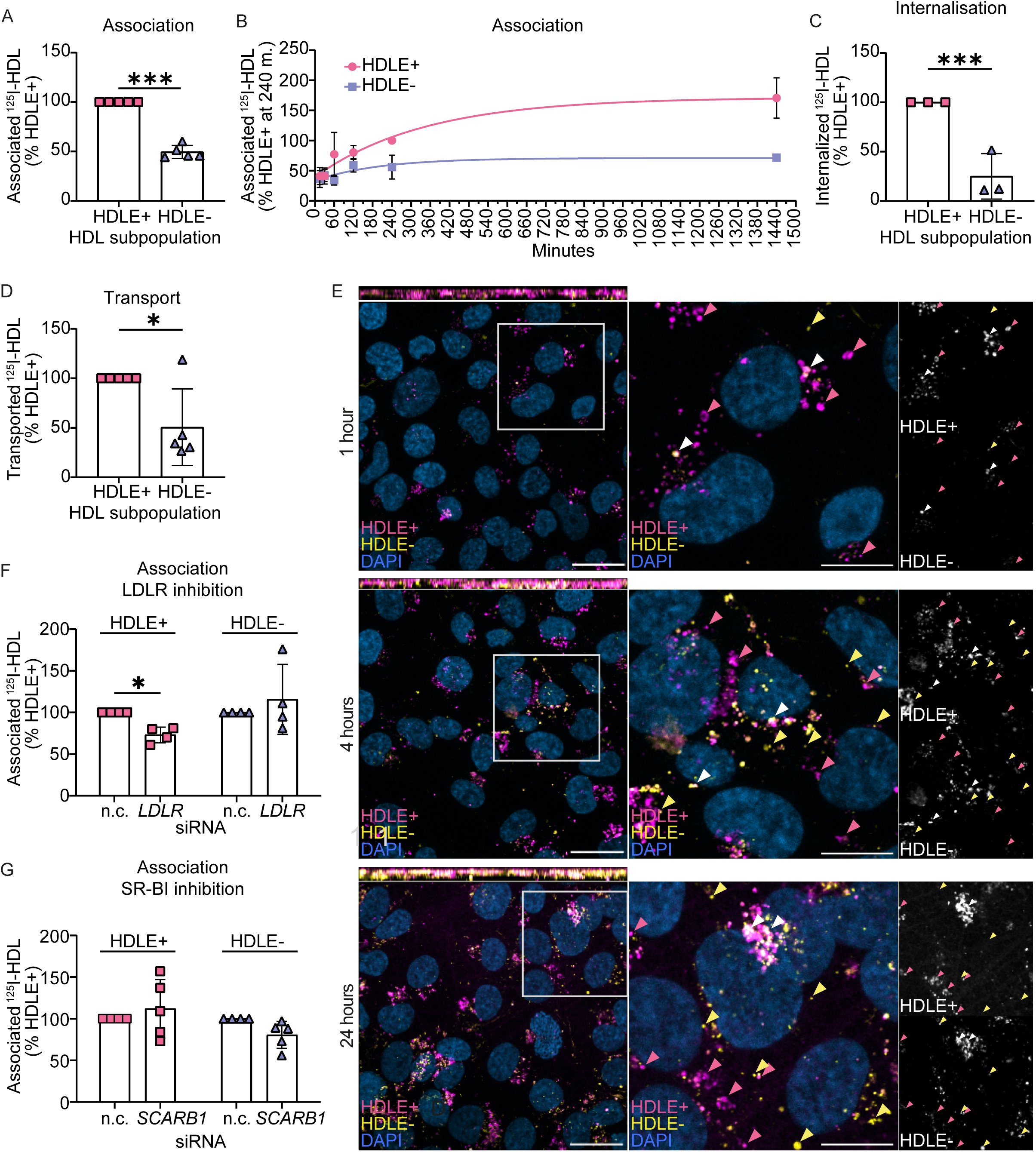
The presence of apoE on HDL increases HDL association, internalization and transport through brain endothelial cells. HDL containing (HDLE+) and lacking (HDLE-) apoE were isolated by immunoaffinity chromatography. ***A)*** ^125^I-HDLE+ and ^125^I-HDLE- associations were measured as described in figure 2 with total HDL as competitor. **B**) ^125^I-HDLE+ and ^125^I-HDLE- association (37°C) was assessed after 15, 30, 60, 120, 240 and 1440 minutes. **C**) Confluent hCMEC/D3 were incubated with Atto655-HDLE+ and Atto488-HDLE-. After 3 hours, cells were washed, detached and median intensity fluorescence (MIF) was measured by flow cytometry. ***E)*** ^125^I-HDLE+ and ^125^I-HDLE transports were measured as described in figure 2 with total HDL as competitor. **F)** hCMEC/D3 were incubated with 50 μg/ml of Atto655- HDLE+ (magenta) and Atto488-HDLE- (yellow) for 1, 4, or 24 hours. Error bar 20 μm in overview and 10 μm in zoom-in. **G**) ^125^I-HDLE+ and ^125^I-HDLE- associations in the absence of LDLR were measured as described in figure 3. **H**) ^125^I-HDLE+ and ^125^I- HDLE- associations in the absence of SR-BI were measured as described in figure 3. Points in graphs represent individual experiments (biological replicates, n = 3-5), bars represent the mean and error bars ± SD, *p<0.05, ***p<0.001.

To further explore the differential trafficking of HDLE+ and HDLE-, we investigated cellular uptake using confocal microscopy. After labeling HDLE+ with Atto655 and HDLE- with Atto488, we found that HDLE+ (magenta) did not fully co-localize with HDLE- (yellow) after 1, 4 and 24 hours (***Figure 4E*)**. We then investigated the role of LDLR and SR-BI in HDLE+ or HDLE- association. Upon silencing of LDLR, the association of ^125^I-HDLE+ but not ^125^I-HDLE- was significantly reduced (***Figure 4F***). HDLE+ internalization was also reduced after silencing of LDLR, but no reduction was observed in the internalization of HDLE- (***Supplemental Figure 4A***). Interestingly, in the absence of SR-BI neither HDLE+ nor HDLE- association was significantly reduced, although HDLE- showed a 20% reduction (***Figure 4G***).

We next investigated HDL binding to brain endothelial cells by cooling down the cells at 4°C to block internalization. First, we measured the specificity of ^125^I-HDL binding using 40-fold excess of unlabeled HDL, LDL or albumin. The addition of HDL or LDL but not albumin significantly reduced ^125^I-HDL binding to hCMEC/D3 suggesting similar receptors for HDL and LDL (***Figure 5A***). Because we previously reported that SR-BI is not required for LDL to bind to brain endothelial cells ^19^, we investigated the role of LDLR. Surprisingly silencing LDLR did not alter ^125^I-HDL binding to brain endothelial cells (***Figure 5B***). Similarly, overexpression of LDLR also did not change ^125^I-HDL binding (***Figure 5C***). Importantly, overexpression of LDLR significantly increased LDL binding confirming LDLR functionality after overexpression (***Supplemental Figure 4B***). These results were confirmed using an antibody blocking the HDL binding site on LDLR, which did not alter ^125^I-HDL association (***Supplemental Figure 4C***). Notably, an antibody blocking the HDL binding site on SR-BI reduced ^125^I-HDL association (***Supplemental Figure 4D***). We then investigated if the presence of apoE on HDL affect HDL binding to brain endothelial cells. Similar to that observed for association and transport, brain endothelial cells bound more HDLE+ than HDLE- (***Figure 5D***). Together, these results suggest that LDLR facilitates the trafficking of HDL independently of direct ligand binding, particularly of HDL particles that contain apoE. To further understand how HDL binds to brain endothelial cells, we silenced the expression of other known apoE receptors namely apoE receptor 2 (apoER2, LRP8) and LDLR-related protein 1 (LRP1), both members of the LDLR superfamily ^34^. The knockdown efficiency of both receptors was confirmed by real-time quantitative PCR (***Supplemental Figure 4E-F***) but their absence did not reduce, but rather slightly enhanced ^125^I-HDL binding to brain endothelial cells (***Figure 5E-F***). Because lipid-free apoE has been reported to bind to heparan sulfate proteoglycan (HSPG) ^35,36^ and we previously found that HDLE+ colocalizes with HSPG ^37^, we next investigated their role in HDL binding to brain endothelial cells. Pre-treatment of hCMEC/D3 with heparin to saturate HSPG binding site on endothelial cells lead to a significant reduction of ^125^I- HDL association (***Figure 5G***).

**Figure 5:**
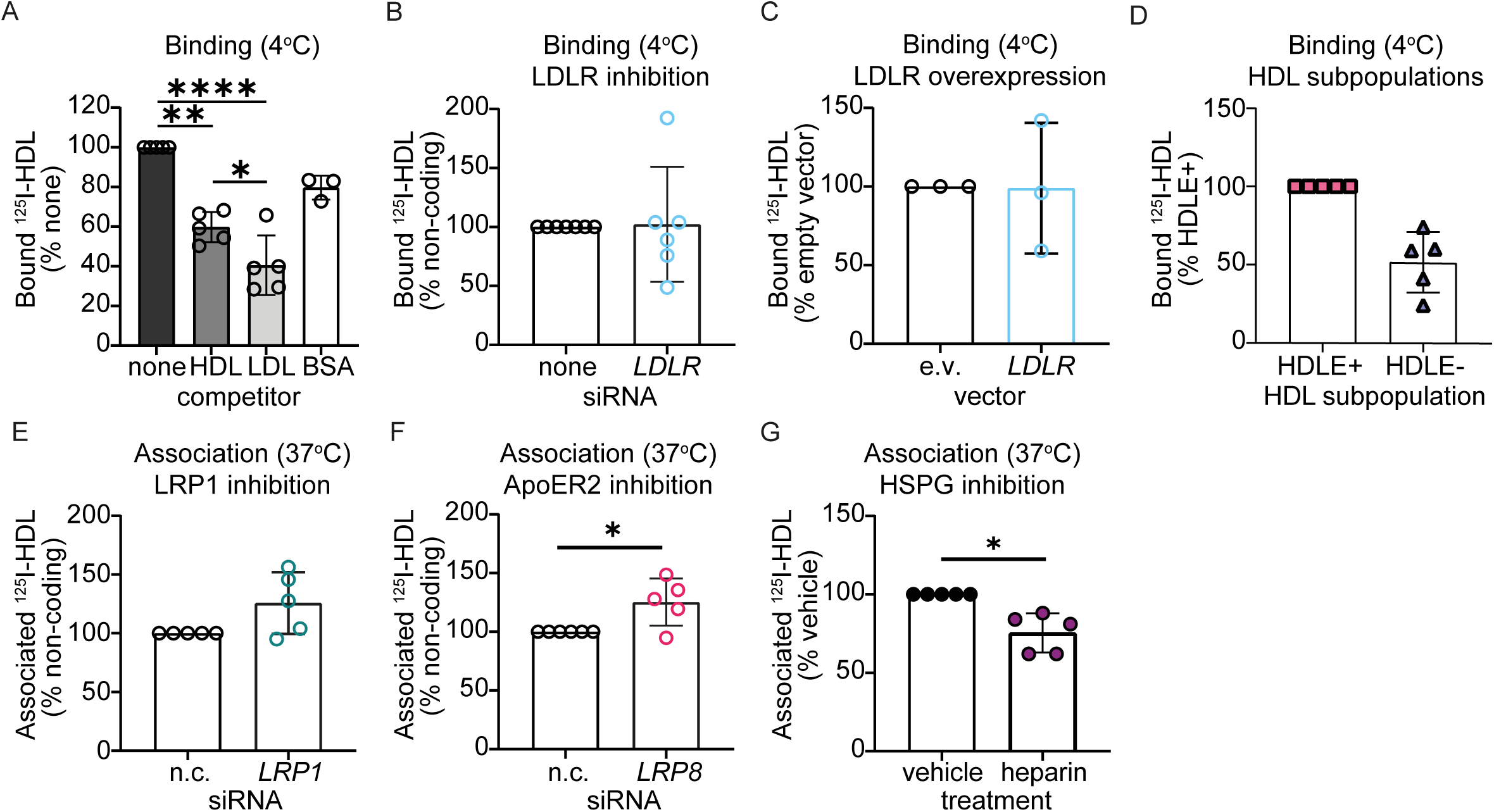
**Heparan sulfate proteoglycans but not LDLR mediate HDL binding to brain endothelial cells**. **A**) Confluent hCMEC/D3 were cooled down to 4°C to block internalization processes. After 30 minutes on ice, cells were incubated with 20 μg/ml of ^125^I-HDL without (none) or with 40× excess of non-labeled HDL, LDL, or BSA as competitors. After 1 hour at 4°C, cells were washed, lysed and radioactivity measured using a γ-counter. **B**) HDLE+ and HDLE- binding with total HDL as competitor were measured as described above. **C-D**)^125^I-HDL binding after RNA interference against LDLR (**C**) and after LDLR transient overexpression (**D**) were measured as above. **E-*F)*** ^125^I-HDL association (37°C) after RNA interference against LRP1 (**E**) or apoER2/LRP8 (**F**) were measured as described in figure 2. **G**) Confluent hCMEC/D3 were incubated with 0.2 mg/ml of heparin. After an hour, ^125^I-HDL association (37°C) was measured as described in figure 2. Points in graphs represent individual experiments (biological replicates, n = 3-5), bars represent the mean and error bars ± SD, *p<0.05, ***p<0.001.

### 4. HDL entry within the CNS is reduced in the absence of LDLR

To assess the relevance of our *in vitro* results, we measured HDL levels in the brain of *Ldlr^-/-^* mice following tail vein injection. Initially, we performed a pilot experiment to estimate the required sample size. Three *Ldlr^-/-^*and three wild type male mice were injected in the tail vein with 100 mg/kg of Atto655-HDL (***Figure 6A***). After four hours and extensive perfusion, HDL level in the brain was quantified using human apoA-I ELISA (***Figure 6B***). Using the average and respective standard deviation, we calculated that 15 animals per group would be sufficient to achieve a power of 80% and α=0.05.

**Figure 6:**
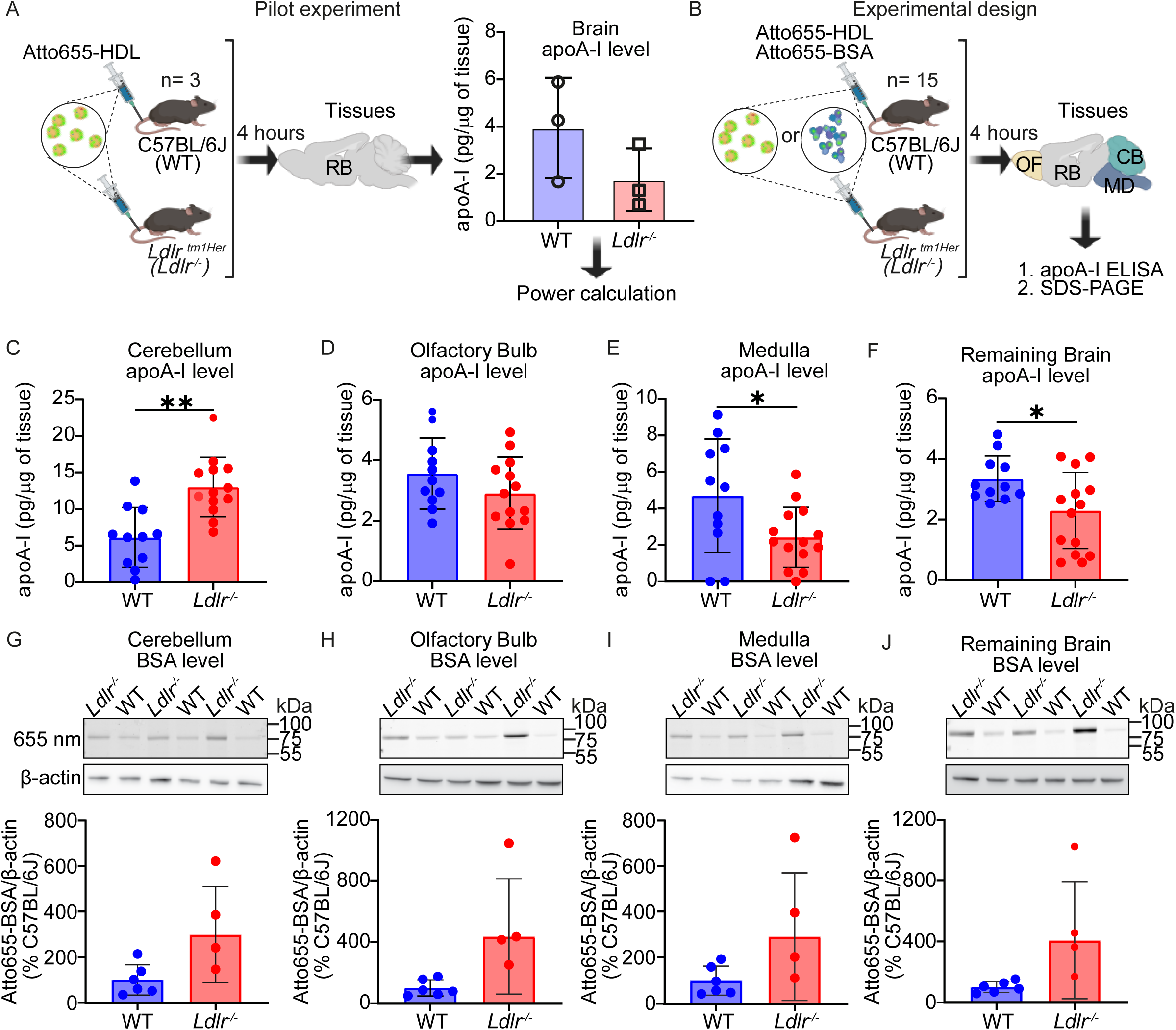
HDL level is reduced in the brain of Ldlr^-/-^ mice except in the olfactory bulb and cerebellum. **A)** Schematic representation of the pilot experiment with Atto655-HDL injected in the tail veins of 3 wild type and 3 Ldlr^-/-^mice to determine sample size for the primary endpoint of brain apoA-I level. ApoA-I level in the brain of Ldlr^-/-^and wild type mice was measured 4 hours after tail vein injection and extensive cardiac perfusion. **B**) Schematic representation of the full experiment, with 15 wild type or Ldlr^-/-^mice were injected in the tail vein. **C-F**) Atto655-HDL (100 mg/kg) were injected in the tail veins of 15 wild type and 15 Ldlr^-/-^ male mice (12-14 weeks). After collection, brains were dissected to isolate the medulla (MD), cerebellum (CB), olfactory bulb (OF) and the remaining brain (RB) tissues. After lysis and homogenization, apoA-I level was measured in 100 μg of total protein from each tissue lysate using a human-specific apoAI ELISA. **G-J**) Atto655-BSA was injected in the tail veins of 5 wild type and 5 Ldlr^- /-^male mice (12-14 weeks old). Brain tissues were dissected and processed as above and Atto655-BSA signal was recoded after 25 μg of lysate were separated by SDS-PAGE. Band intensity was quantified using Image J and normalized to the level of TATA-binding protein. Graphs represent mean ± SD and each point represents an individual animal. * p<0.05, and ** p<0.01.

In the validation experiment, with a separate cohort of 15 animals per genotype (***Figure 6C***), we found that apoA-I levels were significantly enhanced in the cerebellum of *Ldlr^- /-^* mice and unchanged in the olfactory bulb (***Figure 6D-E***) while it was significantly reduced in both the medulla and remaining brain tissues (RB) (***Figure 6F-G***). To understand this observation, we injected Atto655-BSA into the tail veins of both *Ldlr^-/-^* and wild type mice to measure BBB leakage and observed more Atto655-BSA in all regions (***Figure 7H-K***). Together, our results show that despite increased permeability of the BBB in *Ldlr^-/-^* mice, HDL level are significantly reduced in certain brain regions supporting the role of LDLR in HDL transport through the brain endothelium and suggesting that HDL entry in the CNS is region specific and mediated by various receptors.

## Discussion

HDL play pivotal roles in systemic cholesterol transport as well as protection and repair of various tissues ^38^. Despite their hepatic and intestinal origins, HDL and their primary protein component, apolipoprotein A-I (apoA-I), have been identified within the CNS, including the cerebrospinal fluid (CSF), vascular walls, and brain parenchyma ^39^. Moreover, after intravenous injection into animals, native HDL as well as bioengineered HDL-like nanoparticles were recovered in the brain and found to improve the outcome of various experimental disease models including stroke, multiple sclerosis, glioblastoma, and Alzheimer’s disease ^40–50^. These findings suggest an active mechanism enabling HDL translocation across the BBB, a highly selective interface formed by brain endothelial cells. Brain endothelial cells exhibit distinct properties compared to their peripheral counterparts, including specialized tight junctions that restrict paracellular transport and unique receptor expression profiles influencing lipoprotein interactions. For instance, while aortic endothelial cells facilitate LDL transcytosis, brain endothelial cells predominantly degrade LDL, preventing its accumulation within the CNS ^19^. Understanding the molecular mechanisms governing HDL transport through brain endothelial cells is crucial, given HDL’s potential neuroprotective functions.

Previous *in vitro* and *in vivo* studies have demonstrated that lipid-free apoA-I enters the CNS via endocytosis by both choroid plexus epithelial cells and brain microvascular endothelial cells. However, despite its continuous generation through HDL particle remodeling by cholesteryl ester transfer protein (CETP) and phospholipid transfer protein (PLTP) ^51,52^, lipid-free apoA-I is rarely detected in circulation, and the pathways facilitating the translocation of lipid-bound HDL remain poorly defined. Brain endothelial cells express multiple receptors implicated in lipoprotein metabolism, including SR-BI and ABCG1 ^15,53–55^. Previous studies have shown that SR-BI regulates HDL trafficking and function in brain endothelial cells ^37,44,56^, a finding we confirm here. In contrast, the absence of ABCG1 did not affect HDL trafficking, further highlighting differences in lipoprotein metabolism between brain and peripheral endothelial cells.

While SR-BI plays a key role in HDL transport, its silencing does not completely abolish HDL uptake, indicating the involvement of additional receptors. Here, we identify LDLR as a critical regulator of HDL transcytosis through brain endothelial cells. Notably, HDL transport was reduced by 75% *in vitro* in the absence of LDLR, specifically impacting the transcytosis of HDLE+. Traditionally, LDLR is associated with LDL uptake and degradation ^57^; however, picomolar concentrations of the pro-inflammatory cytokine IL- 1β can induce LDL transport through human aortic endothelial cells depending on LDLR ^58^. Similarly, we demonstrate that LDLR facilitates the transport of HDL through brain endothelial cells despite not directly binding HDL. Notably, LDLR appears to function as a co-receptor specific for the HDLE+ subpopulation. We ruled out the role of other known apoE receptor, namely apoER2 and LRP1 but we identified HSPG as potential binding partners for HDL in brain endothelial cells. These results are in accordance with our previous study showing that only HDLE+ colocalizes with HSPG^59^. LDLR may thus subsequently facilitate endocytosis and transcytosis of HDL bound to HSPG. This mechanism diverges from classical clathrin-mediated LDLR endocytosis of HDLE+ in hepatocytes, which leads to lysosomal degradation ^33,60–62^.

Interestingly, prior studies showed that apoE-containing lipoproteins first bind HSPG in hepatocytes and neurons before uptake via LRP1, leading to degradation ^63^. In contrast, LRP1 deficiency in brain endothelial cells did not reduce HDL association but rather increased it, suggesting an alternative mechanism that warrants further investigation. Notably, HDL transcytosis via the apoE/LDLR pathway occurs with minimal degradation, preserving HDL integrity.

In vivo, intravenously injected HDL accumulated in various brain regions, with the highest concentrations detected in the medulla, olfactory bulb, and cerebellum. Consistent with our findings, discoidal reconstituted HDL (rHDL) composed of apoE and dimyristoylphosphatidylcholine (DMPC) preferentially localize to the medulla and olfactory bulb following intranasal administration ^64^. Previous studies reported that circulating HDL localizes within brain endothelial cells within an hour of injection, suggesting active transport across the BBB ^43^. We corroborate this observation by demonstrating that HDL accumulates in brain endothelial cells in a vesicular pattern, consistent with *in vitro* findings. Interestingly, HDL was more abundant in veins and capillaries than in arteries, suggesting a role for vascular zonation in CNS HDL transport. Analysis of publicly available single-cell RNA sequencing (scRNA-seq) data revealed that, while *Ldlr* expression is sporadic across endothelial cells of different vascular beds in mice, *Scarb1* is more prominently expressed in veins and capillaries ^65^. Additionally, LDLR deficiency did not reduce apoA-I levels in the olfactory bulb and cerebellum but did so in the medulla and remaining brain regions. This discrepancy may be attributed to increased albumin levels in the different brain regions of *Ldlr-/-* mice, which could mask specific HDL transport pathways by increasing vascular permeability, a phenomenon previously reported ^66–68^. Alternatively, it may reflect regional variations in HDL receptor expression within different brain endothelial populations. For example, *LDLR* is highly expressed in the medulla in murine and human brains ^69^, however it could not be solely attributed to its expression by brain endothelial cells as it is also expressed by other cells such as astrocytes ^70^ and neurons^71,72^. In human scRNAseq of the entorhinal cortex, prefrontal cortex, inferior temporal gyrus, V1 and V2 did not report any difference in *LDLR* and *SCARB1* expression in endothelial cells ^73^. Finally, spatial single nuclei analysis of the human brain shows that endothelial cells in the hypothalamus have the highest expression of LDLR while the expression is lower in the cerebellum ^74^. The expression of *SCARB1* is more uniform among the investigated regions even though some regions such as the olfactory bulb are not reported. Further research is therefore needed to fully elucidate the roles of LDLR and SR-BI in HDL transcytosis. *In vitro* models provide crucial insights into HDL transport as a holoparticle, but *in vivo* studies are essential to confirm these findings and investigate HDL fate post-transcytosis, including potential remodeling or functional alterations within the CNS. Additionally, exploring the impact of different apoE isoforms on HDL transport could yield valuable insights, given their differential effects on cell-surface receptor interactions, lipid metabolism and neurodegeneration ^37^.

In conclusion, HDL transport across brain endothelial cells involves a complex interplay between SR-BI, LDLR, and HSPG, with regional specificity in the brain likely as a function of vascular zonation. These findings have significant implications for CNS lipid homeostasis and neuroprotection. Advancing our understanding of these pathways could pave the way for novel therapeutic strategies targeting neurodegenerative diseases and enhancing CNS drug delivery. HDL is well-recognized for its vasoprotective effects, including interactions with astrocytes and microglia ^37,48,59,75–77^. Understanding the mechanisms underlying HDL translocation across the BBB could inform targeted therapies leveraging HDL-mediated neuroprotection and facilitate the development of HDL-based delivery systems for CNS-directed treatments, particularly if specific HDL subpopulations can be tailored for distinct brain regions.

## Authors Contributions

Conceptualization and study design: For in vitro work S.K., A.v.E. and J.R, For in vivo work S.K. M.G., W.C. D.V, A.K., V.K., T.B., E.C., C.W. and J.R.; Conducting in vivo experiments: S.K., M.G. W.C, I.D.G, J.F, C.B.; Conducting in vitro experiments: S.K., S.B., E.S., M.C., E.V., N.J., J.R.; Data analysis and interpretation: S.K. M.G., W.C, I.D.G, J.R; Writing—original draft preparation, S.K., and J.R.; Writing—review and editing, S.K., M.G. W.C, I.D.G, D.V., S.B., E.S., M.C., E.V., J.F., C.B., N.J., E.C., C.W., A.v.E., J.R.; Supervision and funding acquisition: J.R., A.v.E., C.W., and E.C.

## Acknowledgements

JR was supported by grants from the BrightFocus Foundation (No. A2021037S) and the Synapsis Foundation Switzerland for Dementia Research (No. 2022-PI04), the Lipidology price from the German, Austrian and Swiss (D-A-CH) Gesellschaft Prävention von Herz-Kreislauf-Erkrankung e.V. and the Swiss Lipid award from the Swiss Atherosclerosis Association (AGLA). AvE was supported by a grant from the Swiss National Science Foundation (No. 185109). JR and AvE were jointly supported by a grand from the Swiss Heart Foundation (No. FF22028). CW was supported by grants from the BrightFocus Foundation (A2021045S) and Heart and Stroke Foundation of Canada (05315516). EC was supported by the French National Research Agency (ANR-21-CE17-0023-01) and the Université Paris Cité (Emergence/VASCAGE). The experimental schematics in figures were made using Biorender.com.

---

**Supplementary Figure 1:**
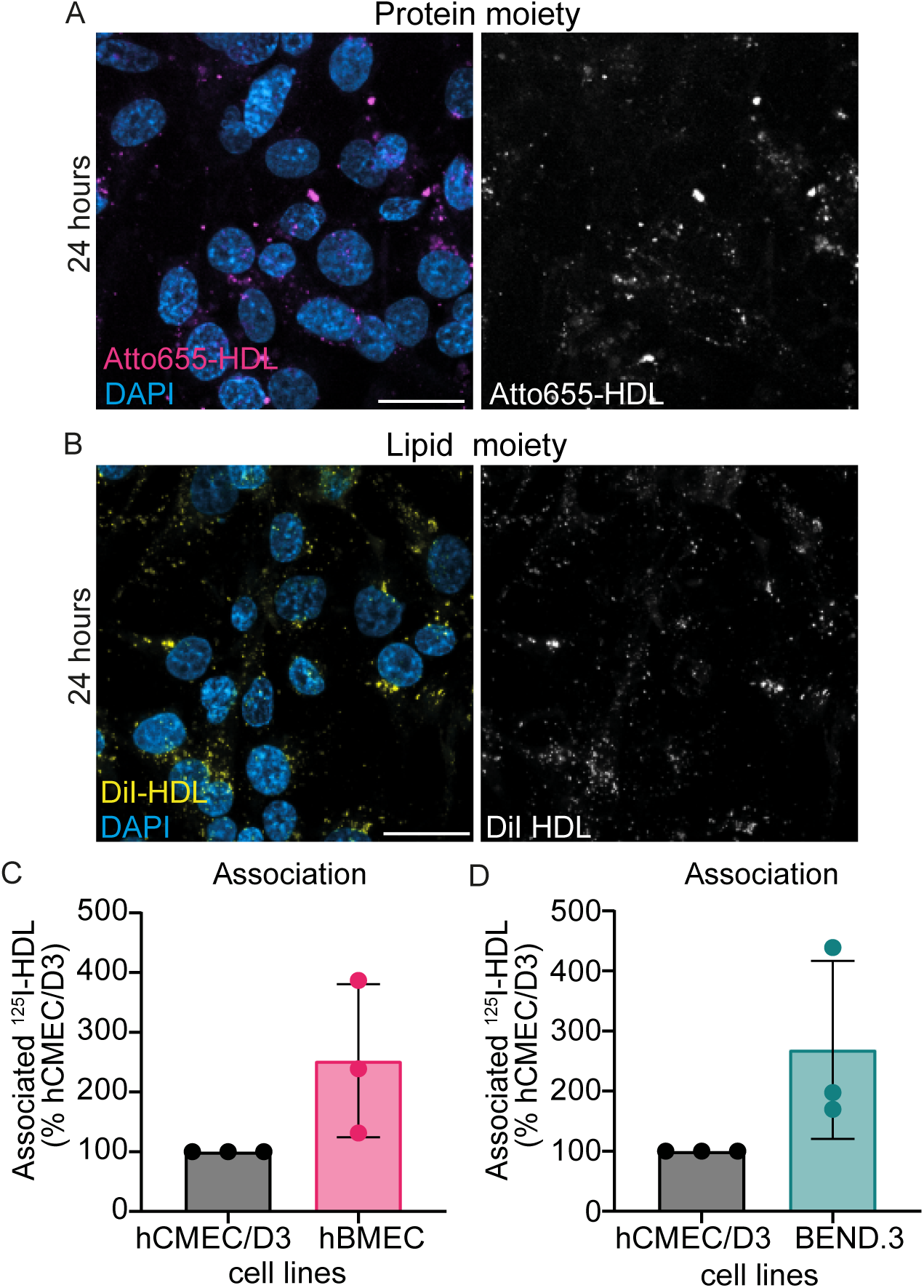
HDL association is similar between the human primary HBMEC and hCMEC/D3 and between HCMEC/D3 and murine cell line BEND.3 A**-B)** Confluent hCMEC/D3 were incubated with Atto655-HDL (A, protein moiety) or DiI- HDL (B, lipid moiety) for 24 hours before counterstaining with DAPI and confocal imaging. **C**) ^125^I-HDL association (37°C) was measured in the brain endothelial cell line hCMEC/D3 and primary human brain microvascular endothelial cells (hBMEC) as described in figure 2. **D**) ^125^I-HDL association (37°C) was measured in hCMEC/D3 and the mouse brain endothelial cell line BEND.3. Points in graphs represent individual experiments (biological replicates, n = 3), bars represent the mean and error bars ± SD,

**Supplementary Figure 2:**
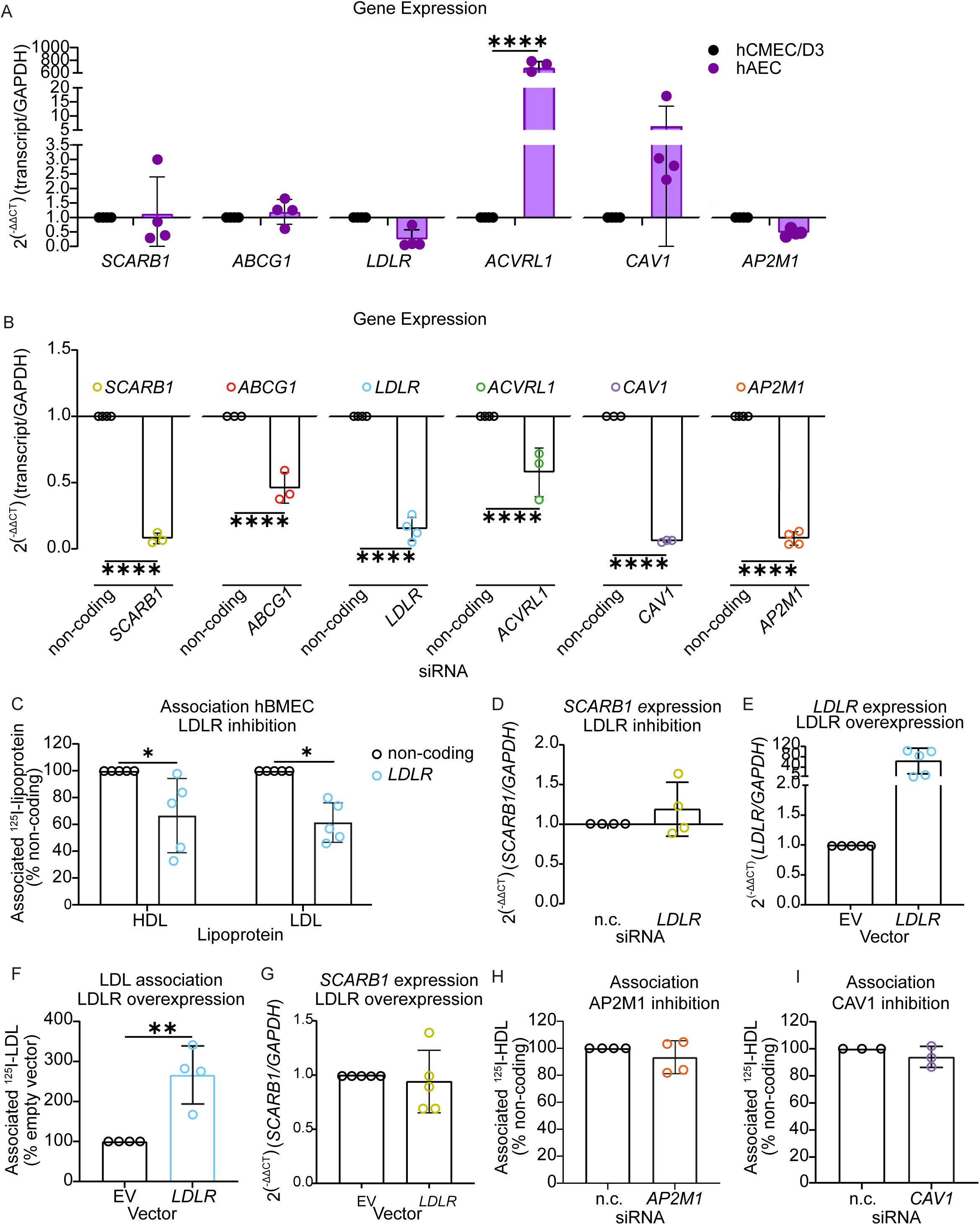
LDLR regulates HDL trafficking in primary human brain microvascular endothelial cells. **A**) Transcript expression of HDL and LDL receptors were measured in hCMEC/D3 and human aortic endothelial cells (hAEC) using RT quantitative PCR. **B**) Transcript expressions after the corresponding RNA interference was measured 72 hours after transfection. **C**) ^125^I-HDL and ^125^I-LDL associations (37°C) in hBMEC lacking LDLR were measured as described in figure 3. (**D**) SCARB1 transcript expression was measured in hCMEC/d3 after RNA silencing against LDLR as above. **E**) LDLR and SCARB1 transcript expression in hCMEC/D3 72 hours after transient overexpression of LDLR. **F**) ^125^I-LDL association was measured in hCMEC/D3 after transient overexpression of LDLR as described in figure 3. **G)** ^125^I- HDL associations in the absence of CAV1 was measured as described above. Points in graphs represent individual experiments (biological replicates, n = 3-5), bars represent the mean and error bars ± SD, *p<0.05, **p<0.01.

**Supplementary Figure 3:**
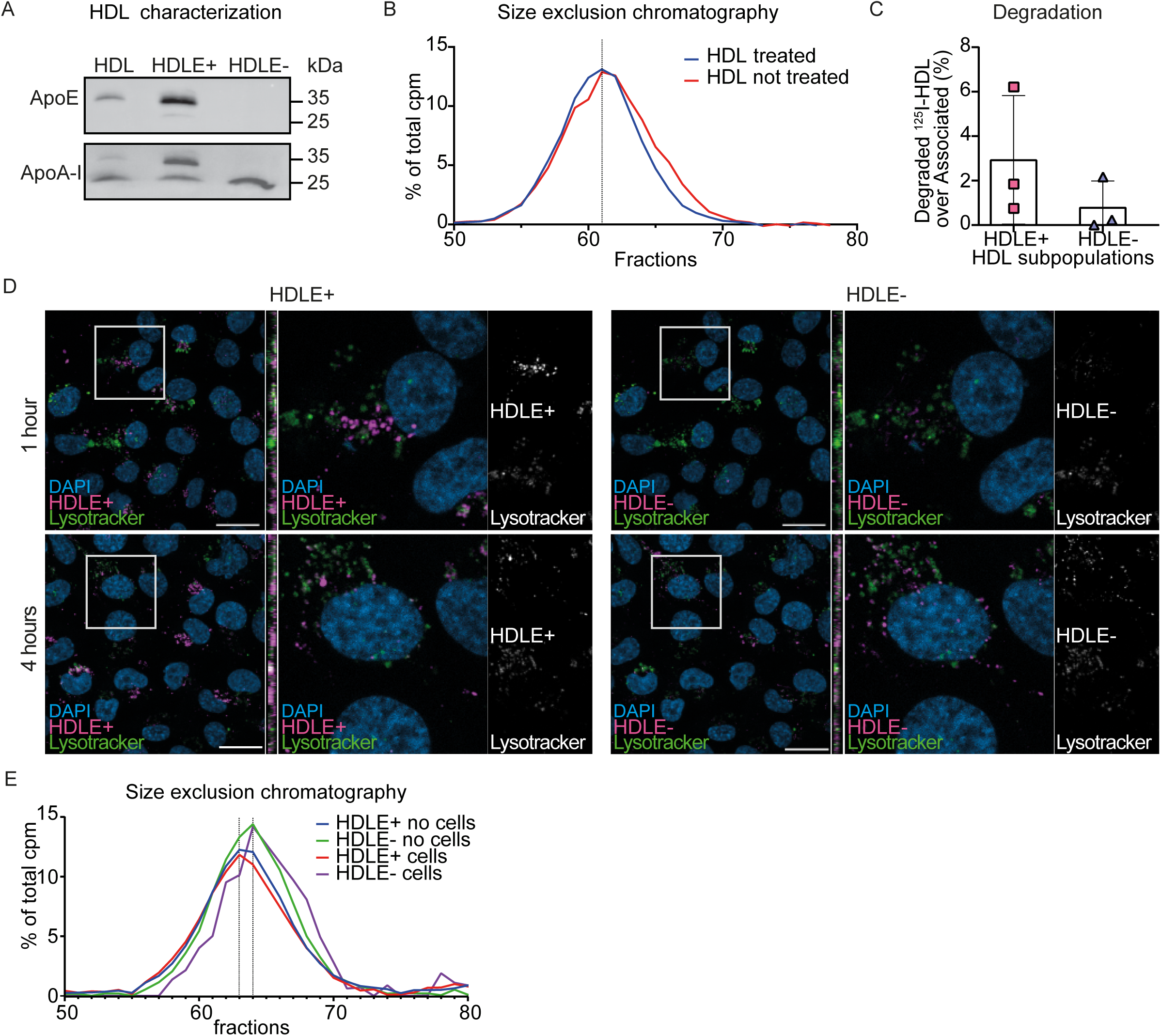
HDLE+ and HDLE- are not degraded by brain endothelial cells. **A**) HDL, HDLE+ or HDLE- (25 μg of total protein) were separated by SDS-PAGE and transferred to PVDF membrane. Expression of apoE followed by apoA-I were probed by Western blotting. **B**) ^125^I-HDL treated with NaSCN at the same ratio as for HDLE+ elution was analyzed by size exclusion chromatography and compared to native ^125^I-HDL. **C**) ^125^I-HDLE+ and ^125^I-HDLE- degradation over association were measured as described in figure 3. **D**) hCMEC/D3 were incubated with 50 μg/ml of Atto655-HDLE+ or Atto488-HDLE- for 1 and 4 hours. After the indicated time, cells were incubated for 15 minutes with 50 nM of lysotracker, washed, fixed, counterstained with DAPI and imaged using confocal microscopy. Bar: 20 μm. **E**) ^125^I-HDLE+ and ^125^I-HDLE- before and after transport were analyzed by size exclusion chromatography as described in figure 2 (representative of 2 individual replicates). Points in graphs represent individual experiments (biological replicates, n = 3), bars represent the mean and error bars ± SD, *<0.05, ***p<0.001.

**Supplementary Figure 4.**
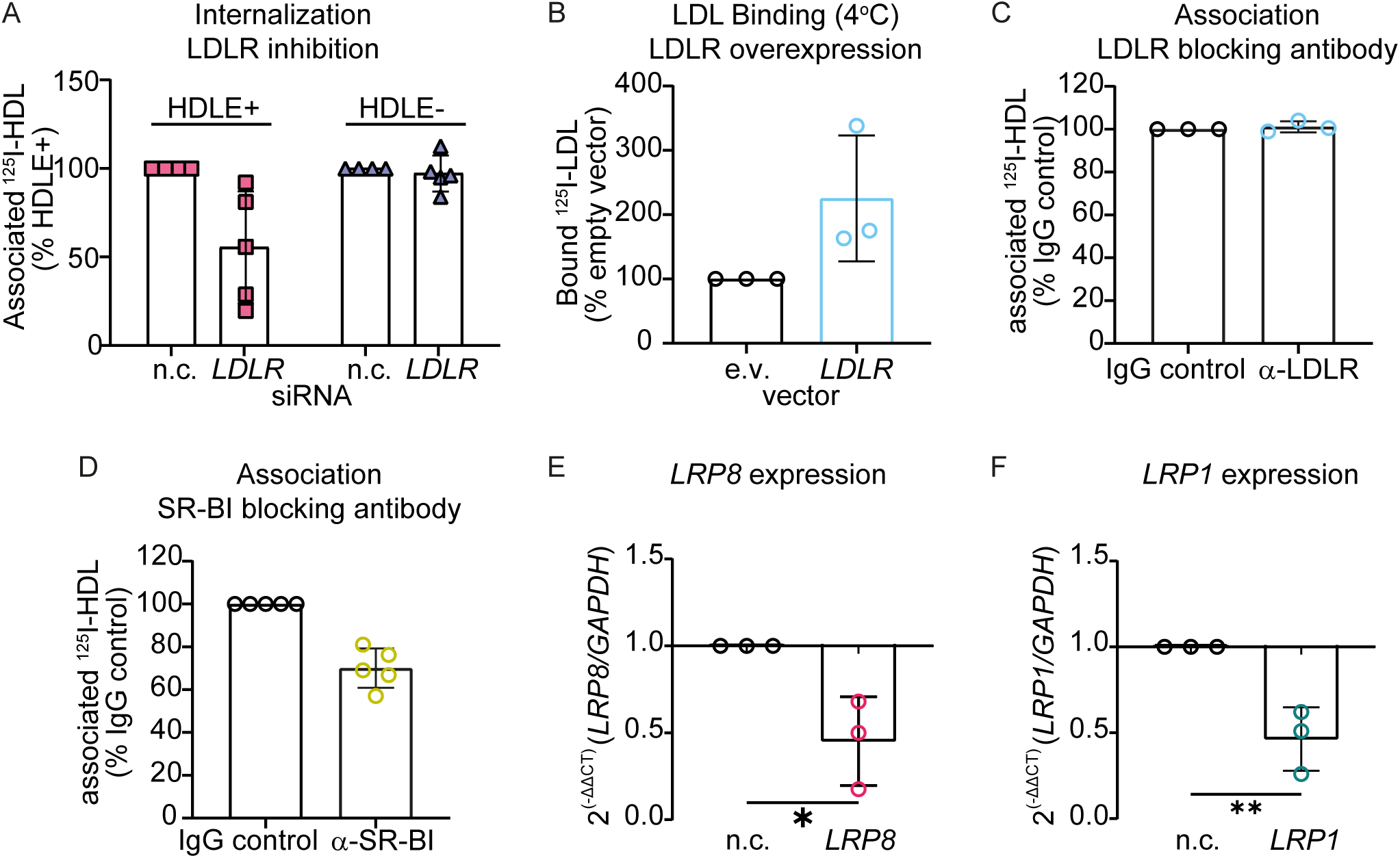
Blocking antibody toSR-BI but not LDLR reduces HDL binding to brain endothelial cells: **A**) Atto655-HDLE+ and Atto488-HDLE- uptakes in the absence of LDLR was measured using flow cytometry as described in figure 3. **B**) Binding of ^125^I-LDL (4°C) was measured 3 days after transient transfection of LDLR. **C-D**) Confluent hCMEC/D3 were incubated with LDLR (B) or SR-BI (C) blocking antibody for 30 minutes prior measuring ^125^I-HDL association (37°C) as described in figure 2. **F-G**) Transcript expressions after the corresponding RNA interference was measured 72 hours after the transfection using quantitative RT PCR. Points in graphs represent individual experiments (biological replicates, n = 3), bars represent the mean and error bars ± SD,

**Supplementary Figure 5.**
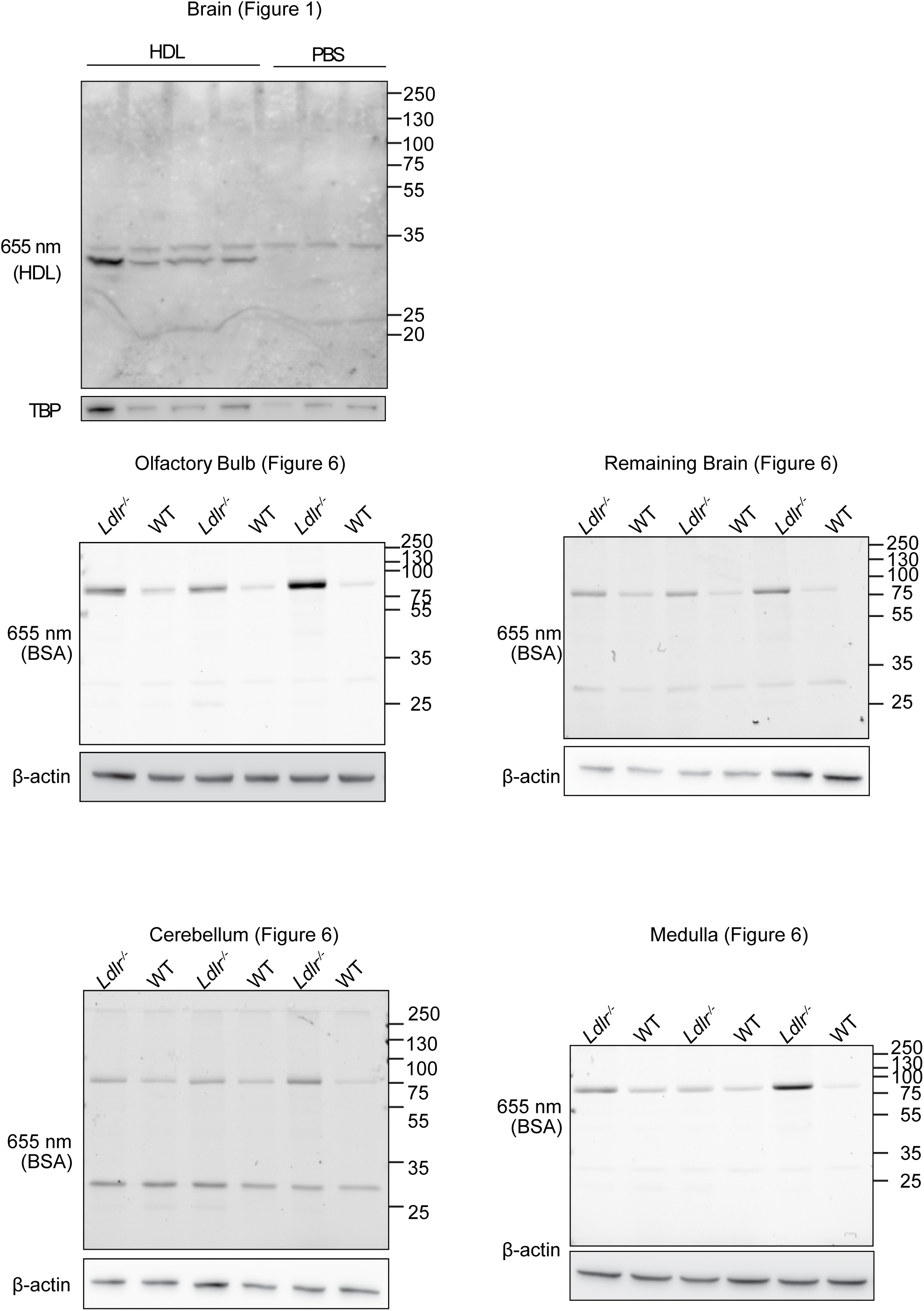
Full blots used in. figures 1 **and 6.** The original and uncropped blot of figure 1 and figure 6.

